# Runx3 Acts as Homodimeric Chromatin Binding Factor Regulating Heterochromatin-Mediated Cancerous Phenotype

**DOI:** 10.1101/2024.08.16.608297

**Authors:** Salma Awad Merghani, Isabelle Bonne, Aik Yong Sim, Gehan Labib Abuelenain

**Affiliations:** Cancer Science Institute, Yong Loo Lin School of Medicine, National University of Singapore; Electron Microscopy Unit, Yong Loo Lin School of Medicine, National University of Singapore; Department of Microbiology and Immunology, Immunology Translational Research Programme, Yong Loo Lin School of Medicine, National University of Singapore; Life Sciences Institute, Immunology Programme, National University of Singapore; Theodor Bilharz Research Institute, Giza, Egypt

**Keywords:** Runx3, Heterochromatin, Homodimer, Nucleosome Binding, Gastric Cancer Running Title: A Novel Regulatory Role of Runx3 in Driving Chromatin Modulations

## Abstract

The recent paradoxical dual citizenships of Runt-related transcription factor 3 (Runx3) in tumorigenesis remains poorly characterized. Here, we report the oncogenic capacity of Runx3 as chromatin modulator in metastatic gastric cancer model. Runx3 exists as homodimer and binds cooperatively to modified nucleosomes. Additionally, we detected a synergistic functional enhancement of octamer transfer, nucleosome sliding and stochiometric integrity of SWI/SNF by Runx3. We found that Runx3 depletion increased nucleosomes occupancy and promoted chromatin silencing by heterochromatin condensation and HP1 oligomerization. ATAC-seq analysis revealed differential accessibility per chromosome due to Runx3 null expression with dysregulation of multiple inflammatory response and DNA repair pathways. Mechanistically, these modulations resulted in aberrant DNA damage repair response, which is rescued by RUNX3 overexpression. These findings reveal a new paradigm in Runx3 biology as dynamic chromatin regulatory element vital for the maintenances of cancerous phenotype.

**Significance:** To the best of our knowledge, the present study is the first to explore the role of Runx3 as homodimeic chromatin binding factor and establishes its oncogenic-function as modulator of heterochromatin de-condensation and SWI/SNF chromatin remodeling activities. These emerging features of Runx3 at the epigenetic level imply a promising direction to screen for anti-Runx3 epigenetic drugs “Epi-drugs” in search of novel gastric cancer treatment.

## Introduction

Gastric cancer (GC) remains the fifth most common malignant cancer and a global burden with expected 62% increase by 2040 (1,2). GC is marked by a high mortality-to-incidence ratio, poor prognosis and survival rate in contrast to other commonly diagnosed cancers (1,2). Epigenetic has emerged as one of the key mechanisms implicated in the development and progression of many cancers including Gastric Cancer (3,4). Epigenetic focuses on understanding changes in gene regulation and expression independent of changes in DNA sequence and include key processes of chromatin alterations, DNA methylation, Micro RNA and Prion (5). Alterations in chromatin structure orchestrate the formation of functionally distinct domains where the DNA is either accessible termed “Euchromatin” or compressed known as “Heterochromatin”. These dynamic chromatin environments function as regulatory switch of many essential processes such as transcription, replication and DNA repair (6). Epigenetic control of gene expression is becoming increasingly vital in understanding cellular neoplastic transformation offering new modalities for diagnosis and future therapeutics (4,6).

Over the past few decades, the function of the Runt-related transcription factors (RUNX) family in cancer has been extensively studied and has been reported to act as tumor suppressors (7). Earlier studies showed loss of RUNX3 expression due to hemizygous deletion and DNA hyper-methylation of promoter region in 45-60% of gastric cancer patients (8). Moreover, heterozygous knockout of Runx3 in mouse was shown to induce adenomas in mammary gland, lung and intestine in aged mice (9). These observations supported the notion that Runx3 acts as tumor suppressor particularly at early cancer stages. However, in the last decade the three-members of (RUNX) family has been implicated in the regulation of plethora of oncogenic processes suggesting a dual role of these proteins in different types of cancer (10). For instance, it has been reported that RUNX3 functions as an oncogene in leukemia, basal cell carcinoma, ovarian cancer and pancreatic cancer (11–14). Recently, Runx3 has been implicated in promoting osteosarcomagenesis in human and mouse by inducing the oncogenic Myc expression (15).

Due to their “double agent” feature, RUNX family members are now being considered as promising targets for cancer diagnosis and treatment with Runx3 becoming a potential epigenetic modulator in gastric cancer (7,10, 16). Recently, Runx3 has been shown to specifically essential for enhancing chromatin accessibility to regions highly enriched with IRF, bZIP and PRDM1-like transcription factors motifs during T cell receptor (TCR) stimulation and memory cytotoxic T lymphocyte (CTL) developmental programing (17). Furthermore, mislocalization of the RUNX3 protein to the cytoplasm was shown to affect its tumor suppressor function, which has been reported in different malignancies including gastric cancers (18–21). RUNX3 has been identified as mediator of the Restriction (R) point of cell cycle by recruiting chromatin associated factors such as Trithorax group proteins or Polycomb repressor complexes (22). These findings suggested that Runx3 might be a potential player in regulating chromatin landscape and architecture.

To gain further insights into Runx3 molecular oncogenic switch, we attempted to dissect the anatomy of Runx3 interactions with chromatin. Here, we report for the first time the potential role of Runx3 as a novel chromatin binding factor regulating heterochromatin and displaying activities in a wide range of chromatin remodeling specialized activities crucial for DNA damage repair response. With these new functions we are proposing a new epigenetic mechanism of Runx3 utilizing GC model as a proof of concept.

## Results

### Nuclear RUNX3 localization is associated with the maintenance of the cancerous phenotype in metastatic mesenchymal gastric cancer

Although Runx3 was initially known for its tumor suppression function due to the frequent loss of expression by genetic/epigenetic factors during the early stages of tumorigenesis, this concept is now being challenged with accumulated recent reports showing that RUNX3 is upregulated across the course of tumor development (23). Since Runx3 expression is masked by genetic and epigenetic alterations reported earlier, we attempted to establish the best cellular model to study Runx3 potential oncogenic nature by examining its level in multiple gastric cancer cell lines (Fig.1.A). Our results show that significant elevated expression of Runx3 is associated with poorly differentiated mesenchymal gastric cancerous cell lines HGC-27 followed by MKN45. Since MKN45 has been reported to be moderately metastatic (24), we choose the metastatic HGC-27 to elucidate the mechanism of Runx3 oncogenesis. Next, we generated a CRISPR knockout of RUNX3 (Rx3KO) in HGC-27 cell lines and confirmed the complete depletion of Runx3 protein in two clones (KO1, KO2) by Western Blotting (WB) and confirmed its nuclear localization via immunofluorescence staining (Fig.1.B). In term of morphological changes, no significant differences were observed between Wild Type (WT) HGC-27 cells and Rx3KO in the standard 2D monolayer culture (Fig.1.C). Interestingly, the 3D-cultured cells formed 3D spheroids with extensive contacts between the cells (Fig.1D). Rx3KO cells formed fuzzy irregular spheroids within 24hr that matured into smaller compacted spheroids after 48hr compared to WT (Fig.1.D). These 3D-morphological changes indicated that Runx3 may play an important role in tumor plasticity possibly through higher order intragenomic and chromatin structures consistent with previous reports (25).

**Fig. 1:**
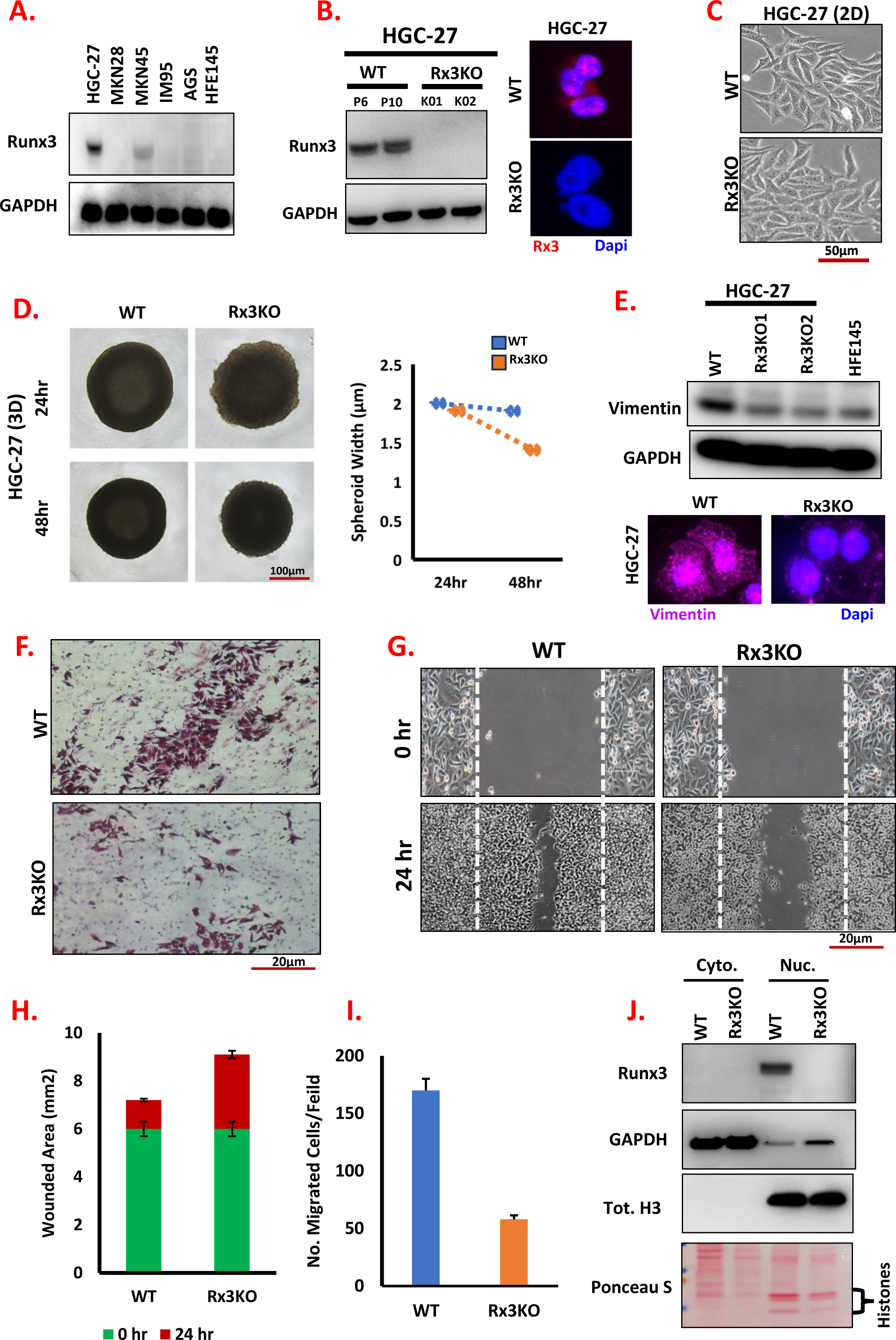
Establishment of Runx3 oncogenic cellular model. **A)**. Evaluation of Runx3 expression in different gastric cancer cell lines including the normal gastric strain HFE145. The mesenchymal poorly-differentiated gastric cancer cell line HGC27 shows the highest level of Runx3 linking it to high grade tumorigenesis. **B)**. CRISPR RUNX3-KO of HGC-27 (Rx3KO) metastatic gastric cell line was created with successful complete loss of Runx3 expression by Western blotting and IF. Example imaging of cell morphology between HGC27 WT and Rx3KO in **C).** 2D monolayer culture and **D).** 3D spheroid formation. Runx3 implicate metastasis in HGC-27 via Reducing: **E)** the expression of Vimentin, **F).** Transwell migration and **G).** Wound Healing scratch, which indicate that WT has higher **H).** chemotaxis and **I)**. cell proliferation rates. **J).** Cellular fractionation shows that Runx3 in the context of HGC-27 has strict nuclear localization.

To explore RUNX3 mediated oncogenesis in HGC-27, we examined several transformed ohenotypes. RUNX3-deficient HGC-27 cell line exhibited downregulation of the metastasis mesenchymal marker vimentin (Fig.1. E) using IF as well as Western Blotting. In addition, the WT possessed higher cell proliferation and invasive rate as seen in the migration (35%) and chemotaxis (77%) abilities which were massively reduced in Rx3KO (Fig.1.F-I). These data indicate that RUNX3 is essential to potentiate the cancerous phenotype and activities in gastric cancer cells.

Recent reports have shown that Runx3 dualistic functions is associated with mislocalization of the protein from the nucleus to the cytoplasm (21). In addition to the IF staining (Fig.1B), we examined the cellular constituent of Runx3 in HGC-27 by isolating cytoplasmic and nuclear fractions, in which nuclear part contains chromatin bound proteins that were released by gentle sonication. We observed that Runx3 strictly localizes in the nuclear/ chromatin fraction with complete absence from the cytoplasm in the WT sample (Fig.1.J, panel 1 lane3). The nil level of Runx3 in the knockout sample confirmed the specific nuclear localisation of oncogenic Runx3, with GAPDH and total H3 demonstrating the clean separation of the two fractions (Fig.1.J, panels 2-4).

### Runx3 Forms Stable Homodimers and Binds to Nucleosomes in a Cooperative Fashion

Given the strong evidence implicating the translocalization of oncogenic Runx3 in the nucleus, we hypothesize the potential function of nuclear Runx3 with chromatin as plausible mechanism for its oncogenic persuasions. To examine the nature of Runx3 association with chromatin, we tested if purified recombinant Runx3 will be able to recognize and bind to chromatin as its substrate similar to other bona fide chromatin factors. To investigate this possibility, we performed quantitative electrophoretic mobility shift assays using purified Runx3-His6 (Fig.S1.A) and fluorescent labelled mononucleosomes (Fig.S1.B) assembled at defined locations on DNA fragments derived from the 601 sequence (26). Titration of Runx3-His6 into reactions containing Cy3-labeled mononucleosomes possessing 36bp of linker DNA on both sides (36W36) resulted in a shifting of the nucleosome signal in concentration-dependent fashion in forms of multiple slower bands (Fig.2.A, lanes 3-4) and is independent of its classic partner CBF-β (Fig. 2.A, lanes 7-8). This shift is diffuse and appears to go through a transition from faster migrating to slower migrating complexes (Fig.2.B, lanes 3-7). However, the extent of Runx3 binding was less when using nucleosomes that had DNA extensions only on one side such as 0W47 or 24W0 (Fig. S1. C-D, indicating the importance of linker DNA on both sides as a perquisite for Runx3-nucleosome binding. As expected, we found that Runx3 was very efficient in DNA binding compared to nucleosomes (Fig.S1.E, lanes 1-3). To decipher the kinetics of Runx3 binding to nucleosomes, we carefully titrated Runx3 and found that the shift is diffuse and appears to go through a transition from faster migrating to slower migrating complexes (Fig.2.B, lanes 3-7 and S1.E, lanes 4-6). A binding curve was constructed by quantifying the shifted nucleosome band. The binding pattern equation is of the form: y = mx + b, with the slope of the regression line coefficient R2=0.849 indicating a strong correlation and association between the protein and its substrate (Fig.2.C). Additionally, we found that Runx3 has higher affinity in binding to nucleosomes assembled using modified histone compared to recombinant nucleosomes (Fig.2.D) and (Fig.S1.F). The pattern of Runx3 binding indicated a cooperative fashion, which suggest that Runx3 might be able to associate with itself in which two or more molecules of Runx3 can bind to a single nucleosome with high affinity.

**Fig.2.**
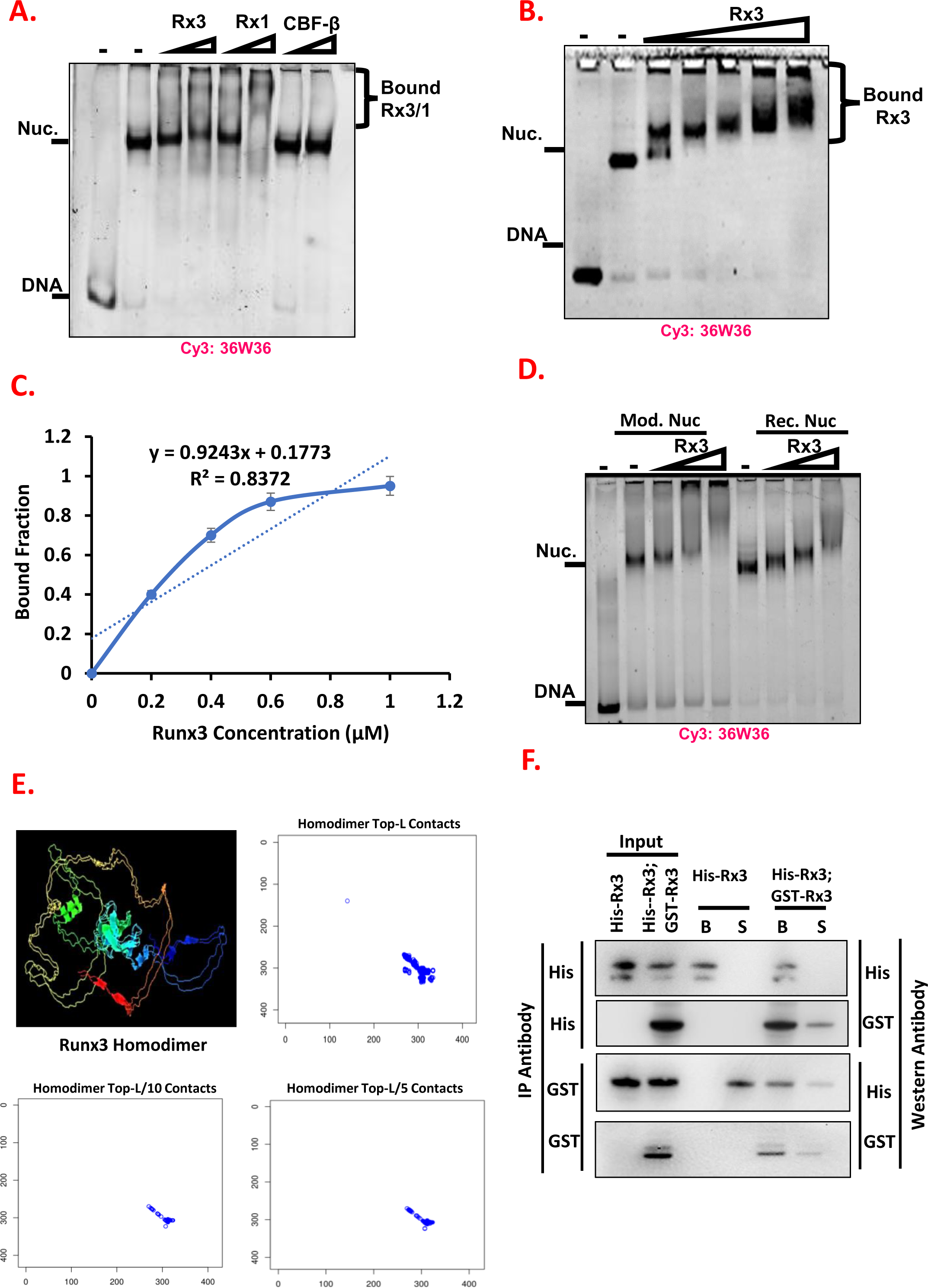
Runx3 binds to chromatin cooperatively and forms stable homodimers. **A**). Binding comparison of Runx3 (Rx3), Runx1 (Rx1) and CBF-β to 30 nm cy3-36W36 Widom template DNA and mononucleosomes. **B).** Incubation of 30 nm 36W36 nucleosomes with increasing concentrations of His-Runx3 (28 nm to 1 μm). **C).** Binding curve of the gel-shifted species (Nuc/Runx3) following incubation with the indicated concentrations of Runx3. **C).** Comparison of Runx3 binding to 30 nm 36W36 nucleosomes assembled with either recombinant octamers (from bacteria) or modified octamer (from chicken). **E).** In silico homodimerization model of Runx3 with the topological contact points using a protein-protein interaction modeling software DeepComplex. **F).** Investigation of Runx3-Runx3 association by performing reciprocal co-immunoprecipitation from a bacteria strain expressing either His-tagged Runx3 (lanes 1, 3-4) or both His- and GST-tagged Fun30 (lanes 2, 5-6). Quantifications were performed for (N=3).

To test if Runx3 can form physiologic homodimers, bacterial strains were made in which Runx3 was expressed as either a His-tagged form only or both His-tagged and GST-tagged forms of Runx3 were co-expressed. Immunoprecipitation of nuclear extracts with the His antibody enriched for His-Runx3 on the beads in both strains (Fig.2.E, first panel). Probing the immunoprecipitants with an antibody against the GST after an IP with the His antibody showed that GST-Runx3 was precipitated with the His antibody only in the co-expressing strain (Fig.2.E, second panel, lane 5). Moreover, the reciprocal immunoprecipitation confirms these findings, following immunoprecipitation with the GST antibody, His-Runx3 was only detected in the co-expressing strain. Additionally, in silico protein-protein docked model of Runx3 generated by DeepComplex (Fig.2.F) using data from NMR alpha fold structures shows high-resolution quaternary structures of Runx3 homodimerers. The homodimers and homomultimers inter-chain contacts were formed at multiple topological levels (Fig.2.E) of the predicted protein L (pentameric/L5 and decametric/L10), which forms mechanical strong interactions (27). These observations indicate that the two differently tagged forms of Runx3 interact in bacteria nuclear extracts, and are in concert with the observation that Runx3 protein prefer to exist predominantly as a homodimer in the nucleus/ chromatin components of the cell.

### RUNX3 Displays Distinct Nucleosome Occupancy Pattern and Regulate Heterochromatin Formation

The strictly nuclear localization and strong nucleosome binding indicated that oncogenic RUNX3 might be involved in chromatin organization and architecture rather than the reported post-translational modifications (7). To test this hypothesis, conventional MNase (micrococcal nuclease) digestion was conducted on wild-type (WT) and Rx3KO nuclei at various time points (Fig.3 A). Interestingly, we observed a significant release of mononucleosomes (fragments size around 147 bp) in WT compared to RX3KO (Fig, 3A, left gel) rapidly at 5min post digestion. This was sustained even after 15min post MNase digestion (Fig. 3.A, right gel), whereas the nucleosome was released slower at Rx3KO clones. This suggested that Runx3 depletion in HGC-27 protected the DNA from MNase digestion by formation of more condensed and higher inaccessible chromatin structure (heterochromatin). Next, we examined the expression level of multiple heterochromatin markers, and we found that deficiency of Runx3 in mesenchymal gastric cancer cells lead to significant upregulation of H3K9 mono-and tri-methylation along with marked reduction of the hallmark of open chromatin H3K4me3 (Fig.3B).

**Fig.3.**
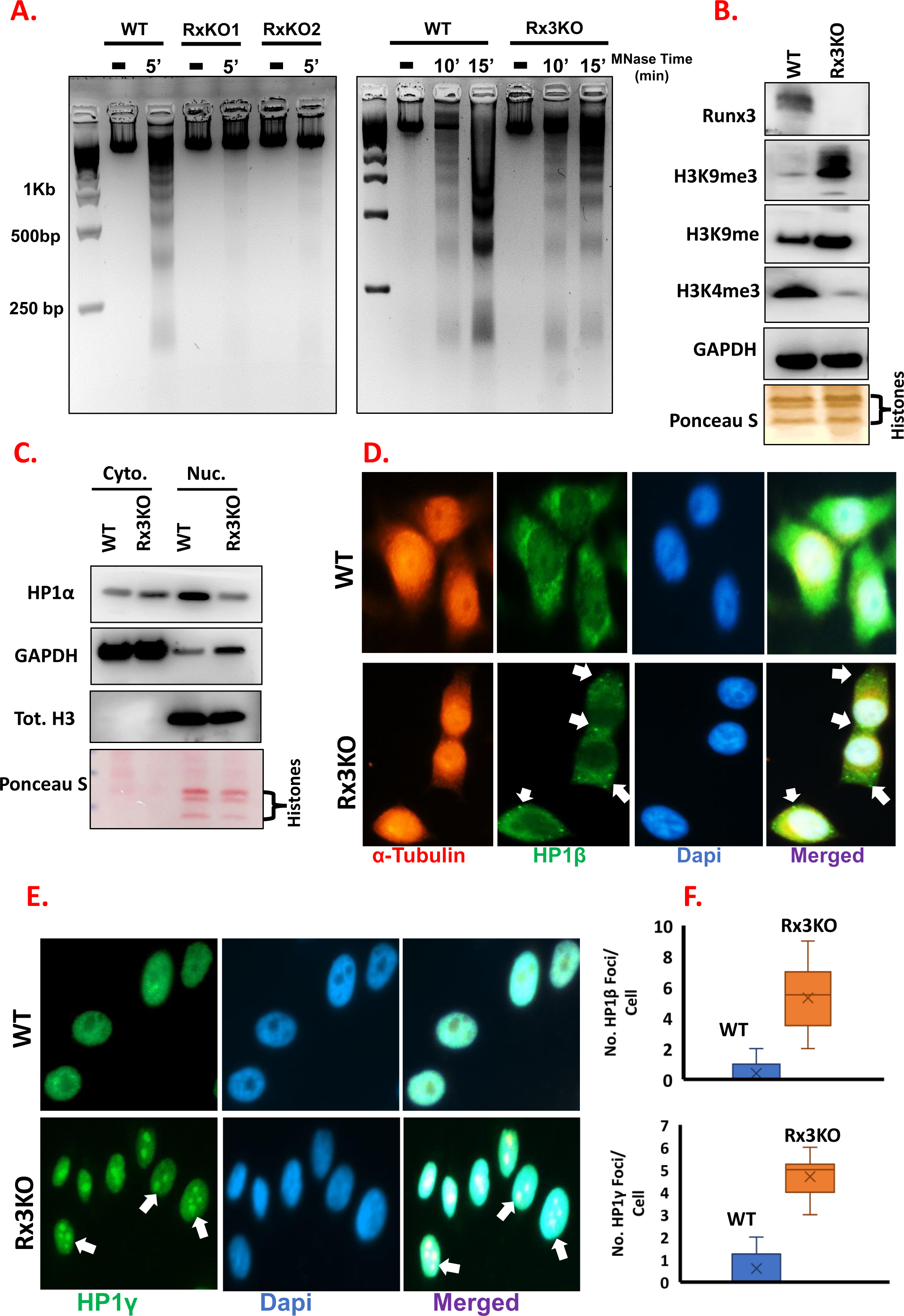
Loss of RUNX3 changes distribution of nucleosome occupancy and heterochromatin dynamics. **A)** MNase digestion profile for HGC-27 WT and Rx3KO nuclei with different concentrations of MNase, followed by nuclei isolation and separation on 3% agarose. **B).** Expression of different heterochromatin/ euchromatin markers using Western Blotting from whole cell extract of HGC-27 WT and Rx3KO. **C).** HP1α localization and level was examined by cytoplasmic/nuclear fractionation. Example of Immunofluorescence (IF) microscopy of HGC-27 WT vs Rx3KO analyzing HP1 isoforms: **D).** HP1β and **E).** HP1γ. **F).** Quantification of Foci formation/ oligomerization between HGC-27 WT and Rx3KO. Quantifications were performed for (N=3).

To further validate this finding, we investigated HP1 (Heterochromatin Protein1) a master player in heterochromatin formation that target H3k9me3 and orchestrate chromatin compaction and silencing (28). Remarkably, Runx3 knockout affected the three isoforms of HP1 differently, it resulted in reduction of HP1α level and its ejection into the cytoplasm (Fig. 3.C). Concurrently, this was associated with massive increase in HP1-β (Fig. 3.D) and -γ (Fig. 3.E) foci numbers with cells containing between 2-5 foci in average in each nucleus (Fig.2. F) with different subcellular localization as HP1γ was strictly nuclear (Fig. 3.D) while HP1β cytoplasmic and at the poles of the cells (Fig. 3.E). These results indicate that Runx3 plays a role in regulating heterochromatin through the distinctive specificities of the HP1 isoforms, in which the decrease in HP1α but not HP1-β and -γ resulted in a global hyper-compaction of chromatin similar to previous studies associating HP1α overexpression with global survival and the occurrence of metastasis (29, 30). These observations contribute to support the novel function of Rux3 in mediating chromatin de-condensation and regulation of heterochromatic foci formation.

### Runx3 Expression is Linked to Heterochromatin Reduction and Periodic Nucleosome Phasing Distribution in HGC-27 Nuclei

In order to gain further insights into the extent of Runx3 mediated epigenetic aberrations in HGC-27, we imaged nuclear compartment in WT and Rx3KO cells using Transmission Electron Microscopy (TEM). In the Rx3KO cells we detected several scattered darkly-stained, irregular aggregations or particles throughout the nucleus or adjacent to the nuclear envelope indicative of highly condensed heterochromatic regions (Fig. 4.A, lower panel). In contrast, the WT cells appeared to have nuclei with no dark patches or ones with less dark aggregations that are mostly single and centrally positioned (Fig. 4.A, upper panel). The absence of heterochromatin dark areas in some of the WT nuclei suggest that the chromatin in these cells is probably the transcriptional active open euchromatin. In the large majority of Rx3KO cells, we observed a high dynamic range of highly condensed nuclei as well as a few brighter euchromatic regions. To compare the average total density of the heterochromatic foci, we examined the fluorescence intensity.

**Fig.4.**
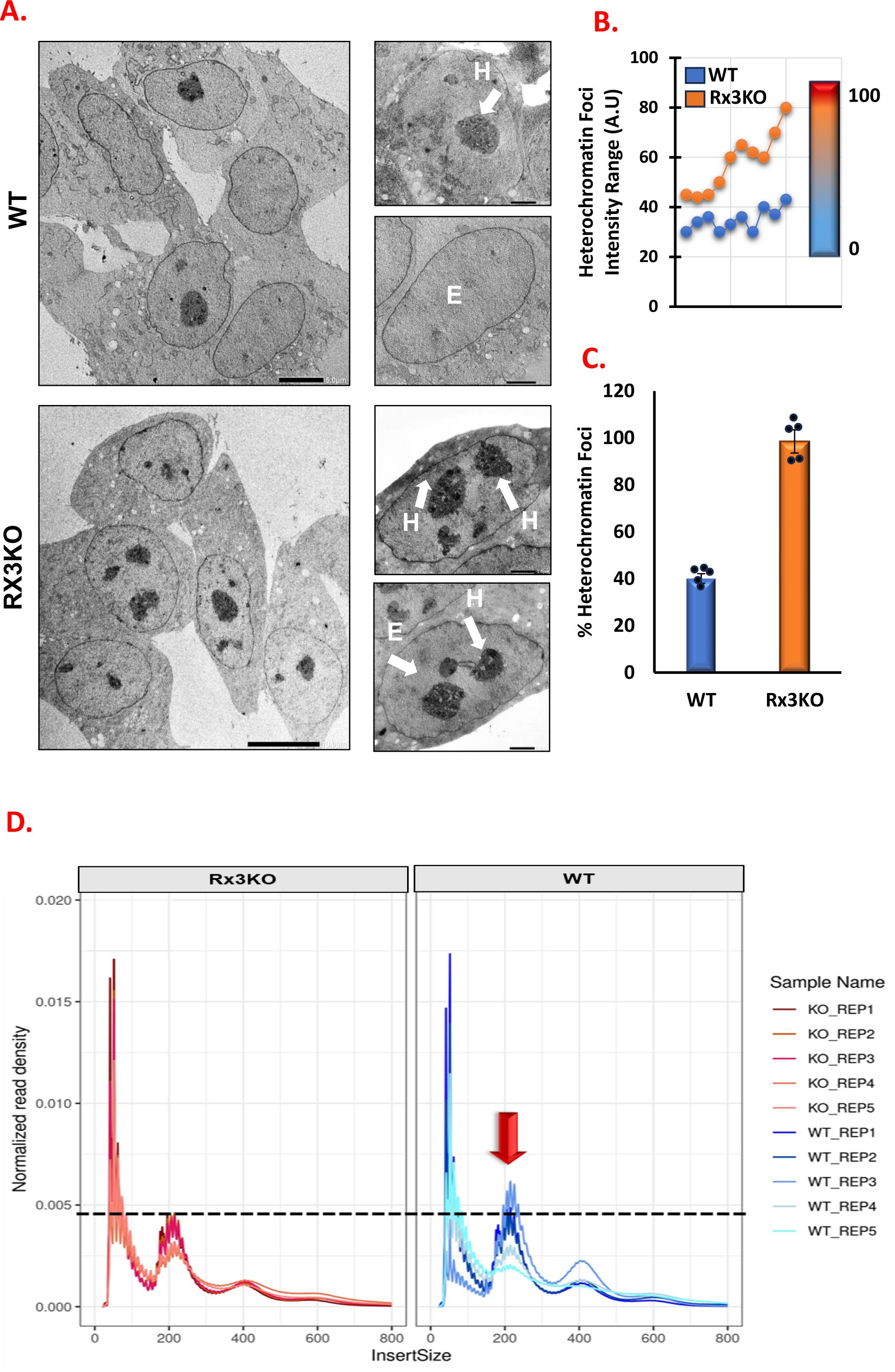
RUNX3 depletion contributes to changes in heterochromatin condensation and nucleosome phasing profile. **A).** Example visualization of Heterochromatin (H) vs Euchromatin (E) by Transmission Electron Microscopy (TEM) in HGC-27 WT and Rx3KO with insets (magnified images) of indicated nuclei (Scale bar: 5 µm). **B).** Quantification of heterochromatin foci average intensity calculated as Arbitrary Unit (A.U). **C).** Average representation of heterochromatic foci in HGC-27 WT vs Rx3KO with individual (Region of Interest) ROI as black dots. **D).** ATAC-Seq nucleosome phasing pattern observed for all samples, with distinct nucleosome-free, mono-nucleosome and di-nucleosome regions. The red arrow specify the mononuckeoosme region/ peak in WT sample while the arbitrary black doted line demarcate the difference in enrichment between WT and Rx3KO samples.

Interestingly, about 90% of Rx3KO nuclei per imaged field expressed condensed chromatin foci (Fig. 4.B) and revealed approximately 2.67-fold increase in the intensity of heterochromatin staining pattern compared to WT (Fig. 4.C). This observation supports the results described earlier, and provide a direct visualization assessment of the effect of RUNX3 deficiency on chromatin architecture inside the nuclei. Since nucleosomes positioning and occupancy studies using MNase titration doesn’t necessarily correlate with chromatin accessibility at promoters, enhancers and other regulatory elements (31), we profiled genomic accessibility using Transposase-accessible Chromatin Sequencing (ATAC-seq) modality. Populations of various fragment lengths ranging from mono, di and trinucleosomes were retained in WT and Rx3KO samples (Fig.4.D). Interestingly, higher density reads of mononucleosomes regions were observed in WT samples (Fig. 4.D, right panel) compared to Rx3KO, consistent with MNase profiling (Fig. 3.A). This clear nucleosome phasing pattern in WT with more prominent periodicity in the overall fragment length distribution pattern is indicative of more efficient Tn5 activity in WT samples which is largely observed in nucleosome-free (NFR) or nucleosome-depleted regions such as mononucleosomes. These data supported regulatory role of Runx3 in chromatin condensation and nucleosomal occupancy.

### RUNX3 Results in Differential Chromosomal Accessibility with Concomitant Dysregulation in several carcinogenic pathways

To further understand the effect of Runx3 on global genome accessibility, we assessed the extent of chromatin accessibility and the magnitude of organizational changes due to RUNX3 deficiency in HGC-27 using ATAC-seq analysis. Differential peaks were identified using DiffBind, the fold-changes of peaks were evaluated, and peaks with a False Discovery Rate (FDR) <= 0.05 were considered differentially accessible (DA). Profile plots over the differentially accessible peaks were also generated from the DiffBind package. We detected an overall genome-wide changes in chromatin accessibility per chromosome as a result of Runx3 modulated expression in HGC-27 cells (Fig.5.A). Interestingly, we observed that in Rx3KO the relatively gene-poor chromosome 18, which is located near the nuclear periphery exhibited more accessibility than in WT samples (Fig.5.A, Ch18). In contracts, WT samples show more accessibility in the similarly sized, gene-dense, and centrally located chromosome 19 (Fig.5.A, Ch19).

**Fig.5.**
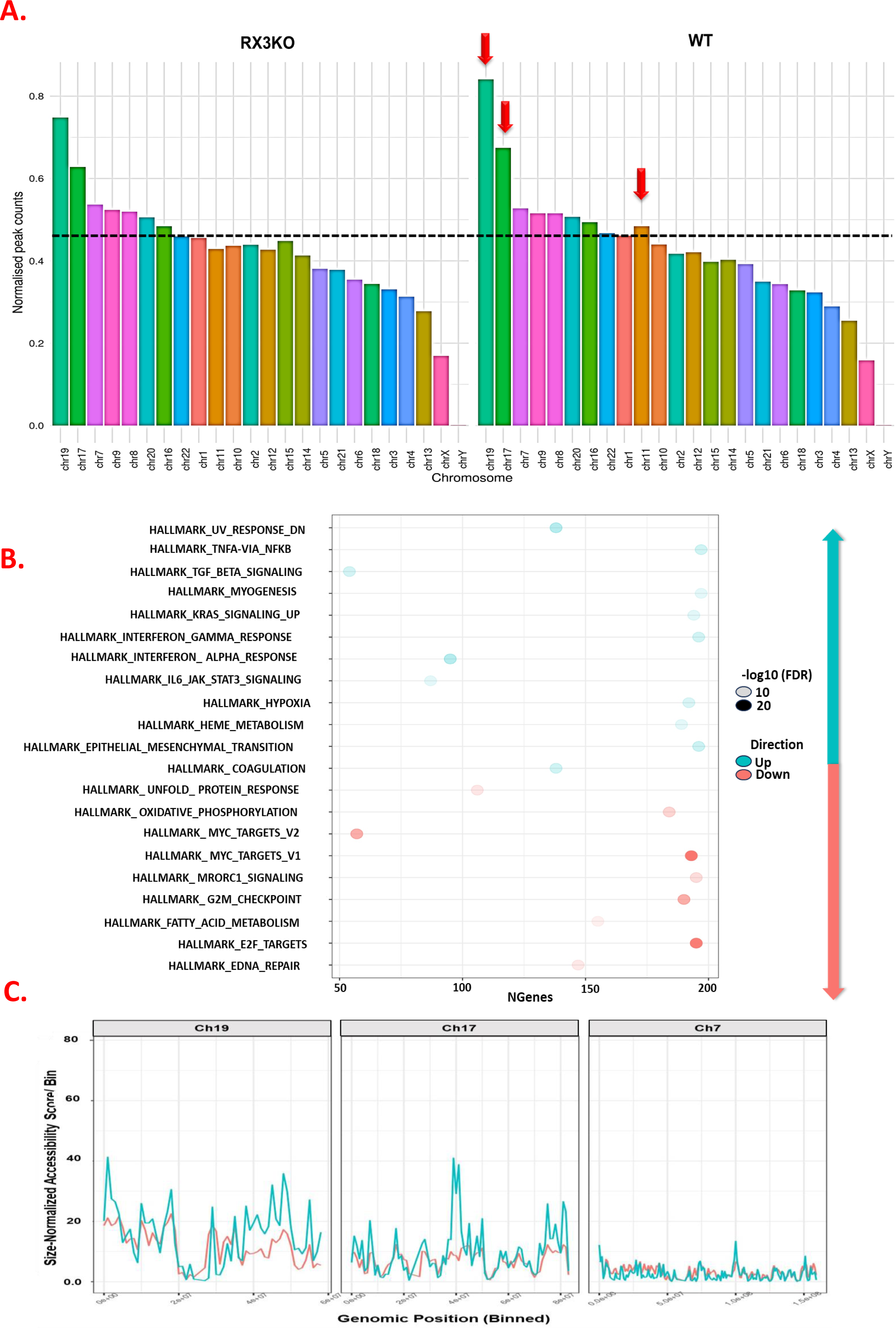
ATAC-seq reveals Runx3-mediated differential chromatin accessibility associated with dysregulation in Myc-targets and DNA repair pathways. **A).** Normalized peak counts of Differentially Accessible (DA) regions in WT vs Rx3KO per chromosome. Most open and accessible chromosomes in WT are indicated by red arrows. Chromosome accessibility score was determined by using the DESEQ2 method and grouping the consensus of ATAC-seq peaks tallied by the number of peaks per chromosome. Followed by normalization of the peak counts by chromosome length that scores between 0 and 1, then scaled these normalised counts by their cumulative sum and a constant scaling factor. B). Integrative RNA-seq and ATAC-seq of enrichment analysis conducted against the ReactomePA pathways and the MSigDB Hallmark gene set in Rx3KO vs WT. C). Representative mapping of chromosome accessibility of open vs closed chromosomes (19, 17 and 7 repectively) in the TCGA STAD patient cohort ATAC-seq Dataset in compassion to our HGC27 ATAC-seq.

We annotated the genomic features of the ATAC-seq peaks with the annotation database from UCSC, interfaced via the TxDb.Hsapiens.UCSC.hg38.knownGene R package. The promoter region is defined to range from -3kb to +3kb of the TSS with both WT and Rx3KO exhibiting a clear enrichment at the TSS. Consistent with this, we also note that a sizeable fraction of the peaks are located within 0-1 kb from the TSS of the nearest gene (Fig.S3.A). As some of the genomic annotations overlap (for instance, a peak may reside within an intergenic region of a gene and also be in the promoter region of another gene), a genomic annotation priority needs to be specified to classify a peak as belonging to one genomic feature or another in the event of an overlap. By plotting stacked horizontal barplots of the genomic annotations across the samples, we observe that the distribution of these annotations are similar across the KO and WT conditions (Fig.S2.B). Using ChIPseeker annotatePeak function, around 70% of the peaks in both WT and Rx3KO reside within either the distal intergenic, intronic or promoter region that is <=1kb from the Transcription Start Site (TSS) (S2.C).

To determine whether changes in open chromatin regions in ATAC-seq analysis were associated with changes in gene expression, we integrated ATAC-seq data with RNA-seq analysis of WT and Rx3KO using the ReactomePA pathways and the MSigDB Hallmark gene set. The enrichment analysis revealed that in Rx3KO Myc-targets, E2F, G2M and DNA repair were among the significant downregulated pathways (Fig. 5.B, Red section), while, inflammatory response pathways such as interferon α/γ and TNF-α were among the top upregulated (Fig. 5.B, Teal section). Immunoblotting analysis of some of these targets showed reduced expression of c-myc along with elevated levels of IFγ (S2.D). These results suggest that loss of RUNX3 leads to negative transcriptional regulation of processes involved in neoplastic cell growth and proliferation. Next, we aligned our ATAC-seq data with The Cancer Genome Atlas Stomach Adenocarcinoma (TCGASTAD) specific peak sets from the GDC ATAC-seq database. Our chromosome accessibility pattern and the genomic annotations overlap results are recapitulated in the TCGASTAD patient cohort ATAC-seq dataset (S2.E-F). We observed strong correlation between the two data sets in which chromosomes 19, 17and 8 are the top most accessible chromosomes (Fig.5.C), whereas chromosomes7 and 9 are the among the least accessible (Fig.5.C).

### Loss of Runx3 Triggers SWI/SNF Complex Dissolution by Stimulating Polyubiquitination and Proteasomal Degradation Pathways

To further explore the impact of newly observed roles of Runx3 in binding directly to chromatin and regulating its compaction, we attempted to study Runx3 interaction network by performing proteomics analysis after Biotin-dependent proximity labelling. We found that Runx3 in the context of Wild Type HGC-27 co-localizes with multiple SNF-2 members in addition to several chromatin organization factors (S3.A). The Gene Ontology (GO) annotations show that the most enriched biological process is the ATP-dependent Chromatin remodeling with a -log10 (p-value) of 3.74e-18 (S3.B). Among the potential interacting partners were multiple subunits of the master regulator and founding member of ATP-dependent chromatin remodeling proteins SWI/SNF (S3.A-B). However, the mass spectrometry identified 1227 candidate proteins and CBF-β the physiologic partner of Runx3 was among stronger group with a >4-fold enrichment and p < 0.01 (S.T1). To further understand the nature of SWI/SNF connection, we examined the expression of SWI/SNF subunits in our WT and Rx3KO cells RUNX3 knockout resulted in marked reduction in the level of SMARCA4/ Brg1 and SMARCA2/ Brm (Fig.6.1, panel 1-2), while ARID1A and 2 were moderately downregulated (Fig.6.1, panel 3-4). The significant decrease in SWI/SNF expression in Rx3KO cells was also detected by indirect immunofluorescence microscopy (Fig.6. B). Differential dysregulation of several chromatin remodeling and modifying complexes was observed at the RNA level with no significant change on SWI/SNF catalytic subunits SMARCA2 and -4 using R2 Profiler Chromatin remodleing and modifications PCR arrays (S4). The observation that Rx3KO cells express significantly less SWI/SNF proteins than WT cells suggested that this reduction may be due to enhanced protein degradation, which is usually induced by ubiquitination. Consequently, we examined the ubiquitination profile in WT and Rx3KO cells, and we found an elevated levels of ubiquitination signal in Rx3KO cells compared to WT specially (Fig.6.C). The enhanced polyubiquitination coincide with increase in the level of L3MBT3 (Histone Methyl-Lysine Binding Protein 3), involved in methylation dependent proteolysis of the SWI/SNF and complex assembly/disassembly (32).

**Fig.6.**
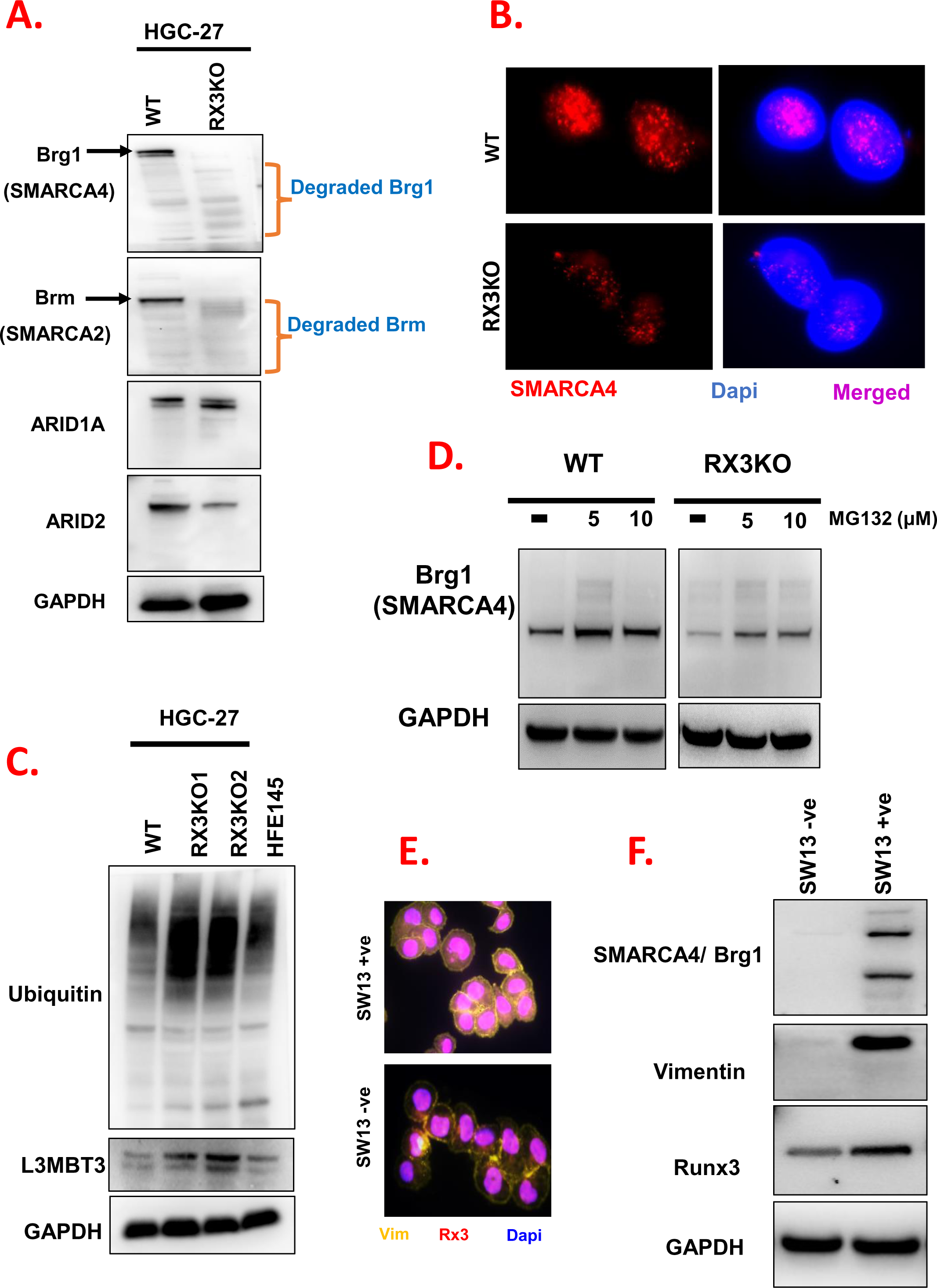
Runx3 regulates the proteolysis of SWI/SNF and play a role in maintaining the complex integrity. **A**). Runx3 ablation downregulates multiple subunits of endogenous mSWI/SNF in HGC-27. Different SWI/SNF subunits were analysed using whole cell lysates of WT and Rx3KO followed by Western Blotting. **B).** Loss of RUNX3 resulted in reduction in the level of the SWI/SNF catalytic subunit SMARCA4 inside the nuclei of Rx3KO by indirect immunofluorescence microscopy. **C).** Increased polyubiquitination and L3MBT3 levels in Rx3KO cells compared to WT by Western analysis of total cell lysates using a specific antibodies. **D).** Recovery of SMARCA4/Brg1 level after treatment with MG-132 proteosome inhibitor. WT and Rx3KO cells were pretreated with MG-132 proteasome inhibitor for the indicated concentrations and were analyzed by immunoblotting after 24hr. Characterization of SW-13 vim-ve and vim +ve subtypes using: **E).** Indirect immunofluorescence microscopy which reveal distinct pattern of vimentin expression, and **F).** Immunoblotting showing unique SMARCA4/Brg1 expression that coincide with vimentin and Runx3 levels.

The accumulation of ubiquitination in the high molecular weight region corresponding to proteins such as SMARCA4 (molecular weight ∼180-200KDa) suggests that Runx3 plays a role in stabilization of SWI/SNF complex and its protection from polyubiquitination which triggers proteasomal degradation. To test whether the proteasome pathway is responsible for SWI/SNF protein breakdown, WT and knockout cells were treated with 5 and 10μM of the specific proteasome inhibitor MG-132, and the catalytic subunit SMARAC4 level was analysed by immunoblotting after 24hr. MG-132 administration resulted in stabilization of SMARCA4 protein and restore its expression in RX3KO cells (Fig. 6.D). These results indicate that Runx3 might be a potential player in maintaining the integrity and stoichiometry of the SWI/SNF complexes controlled by a lysine-methylation dependent ubiquitin proteolysis.

To validate the expression dependency between Runx3 and SWI/SNF expression we utilized the human adenocarcinoma cell line SW13 which is composed of two distinct subtypes, one that expresses neither SMARCA4/BRG1 nor SMARCA2/Brm and is negative for mesenchymal metastasis marker vimentin (SW13(vim−ve)) and the other, which does express both plus vimentin (SW13(vim+ve)). We isolated SW-13 subtypes by FACS sorting using conjugated vimentin antibody (S5.A), and confirmed it by IF staining (Fig.6.E). We found that vimentin expression correlates completely with SWI/SNF subunit SMARCA4 and Runx3 levels in the cell (Fig.6.F). Additionally, the IF staining exhibited similar pattern in which the SW13 vim-ve subtype has lower Runx3 expression (Fig.6.E). These observations suggested a co-regulation and mutual functional association between SWI/SNF and Runx3-medited metastasis.

To further understand the relationship between Runx3 and SMARCA4 in physiological settings. We asked whether Runx3 and SMARCA4 expressions have any clinical significance in gastric cancer patients. To address this, we measured Runx3 level by immunohistochemistry staining in Novus Biologicals-Tissue Array Slide (n=59) of different tumor grades to establish the link between Runx3 and cancer stage (S6.A). We found that low Runx3 expression is observed in well differentiated (Grade I) samples (S6.B), while medium range levels were detected in moderately differentiated (Grade II) tissues (S6.C), and the positive Runx3 expression distinguished the poorly differentiated (Grade III) tumors (S6.D). Next, Kaplan– Meier survival curves of gastric cancer patients m-RNA (n=875) of all TNM tumor stages produced by data obtained from KM-plotter (34) were generated based on the expression levels of Runx3 or/and SMARCAD4 markers (S6.E-F). The results showed that the survival rate of the patients in the Runx3-low group (n=526) was significantly higher (S6.E) than that of the patients in the Runx3-high group (n=349) (P-value=0.0002). Similarly, the survival time of patients with reduced SMARCA4 expression (n=628) was longer (S6.F) than that of patients with high expression (n= 247). The combined probing of Runx3 and SMARCA4 resulted in slight increase in the number of surviving patients (S7.A) with lower expression of both markers (n=546) and even higher survival in Grade III patients (S7.B). Additionally, positive effect as a result of Runx3 or SMARCA4 low expressions was observed in patients with endogenous reduced Runx3 levels (S7.C-D) as well as in the median survival curve (Table.S7.1). Interestingly, stage III patients with low Runx3 levels exhibited an increase in median survival rate of 41.2 months compared to 25.5 months of those with higher Runx3 in the same category (Table.S7.2), and to 36.4 months obtained in all TNM tumor stages (Table.S7.1). These results indicated a positive correlation between the low expression of Runx3 and SMARCA4 with tumor grade and survival rate, especially in the late stages of gastric cancer.

### Runx3 Interacts with SWI/SNF and Enhances its Octamer Transfer and Remodeling Activities

To better understand the nature of interaction between Runx3 and SWI/SNF, we used complementary computational modelling to gain structural and dynamical atomic scale insights on their interactions. The predicted structure based on the available biochemical and biophysical information shows that the Runx3 can form a heterodimer with SMARCA4/Brg1 subunit of the SWI/SNF complex (Fig.7.A). The potential region of interaction is within the evolutionarily conserved Runt domain as the interaction with all SNF-2 member is lost when HGC-27 R122C mutant was subjected to proximity-dependent biotinylation identification (BioID) followed by mass spectrometry (S5.E-F). To validate this interaction in cellular and physiologic settings, we immunoprecipitated (IP) the endogenous SMARCA4/Brg1 of SWI/SNF from HGC-27 WT and Rx3KO cells followed by probing the immunoprecipitants using a specific antibody against Runx3. We detected Runx3 expression and another SWI/SNF SMARCD1 in the WT cells (Fig.7.B, left panels), while the levels of immunoprecipitated SMARCA4 in Rx3KO were severely reduced in Rx3KO cell with moderate expression of SMARCD1 (Fig.7.B, right panels). This indicates that Runx3 is capable of directly interacting with SWI/SNF and maintaining its physiologic expression.

**Fig.7.**
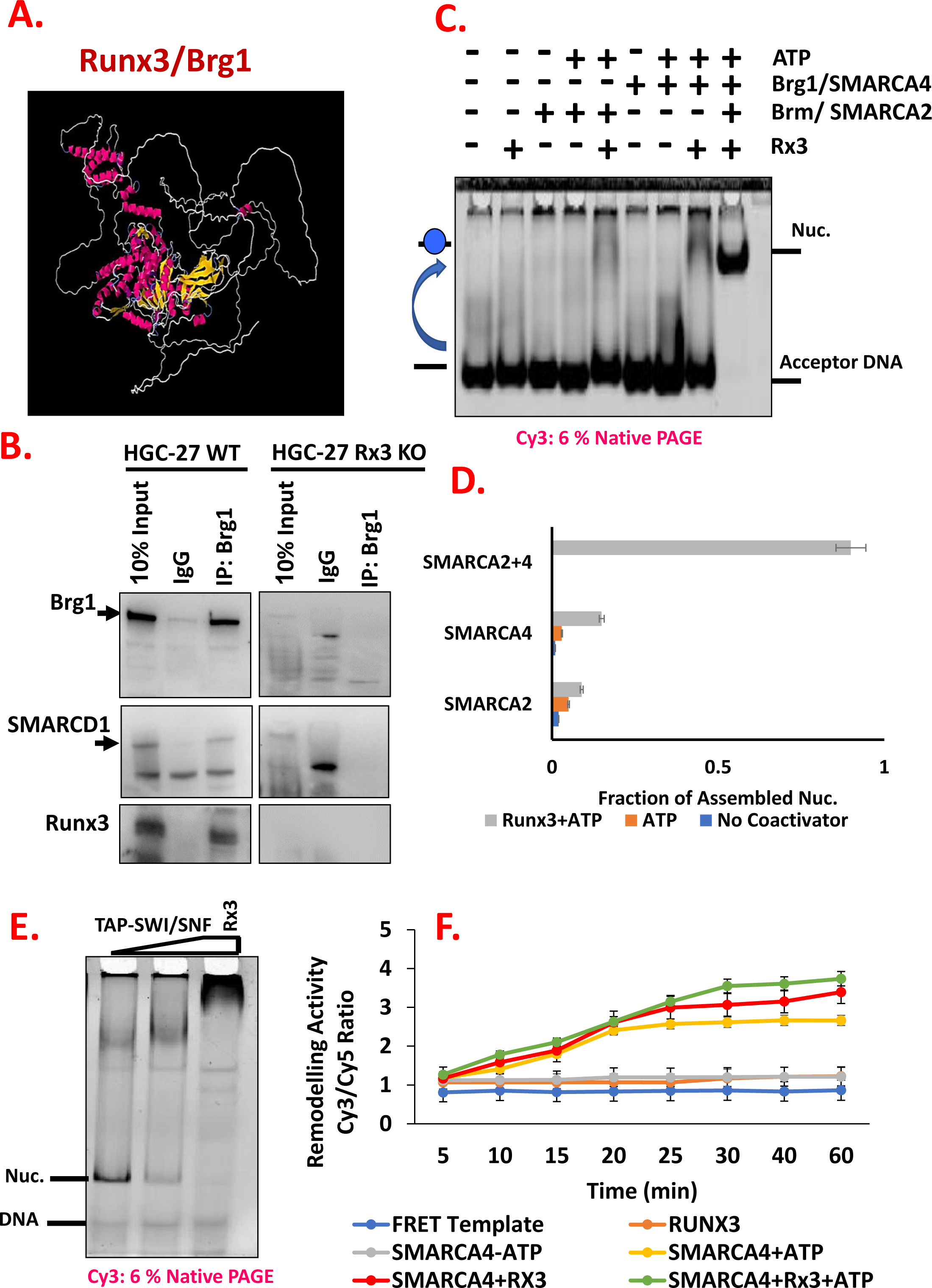
Rux3 interacts with SWI/SNF and synergistically enhances its chromatin remodeling activities. **A).** Inter-chain Contact Prediction and Distance-Based Modelling of Runx3 heterodimerization with SMARCA4 catalytic subunit of SWI/SNF. **B).** Western blots of Runx3, SMARCAD4/Brg1 and SMARCAD1 recovered following immunoprecipitation using specific antibody against Brg1/ SMARCAD4 from HGC-27 WT and Rx3KO. **C).** Octamer-transfer assay examining Runx3 effect on SWI/SNF catalytic subunits nucleosome transfer In-Trans. Briefly, an equal amount of the SMARCAD2/4 subunits were added to 100 ng of the 36W36 DNA template in the presence/absence of 0.8μM of Runx3 and 1 mM ATP for 1.5 h at 30 °C in the presence of 10 ng of donor nucleosomes. The binding reactions were then resolved on a 6% mini-acrylamide native gel at 150 volts for 1 h.. **D).** Quantification of transferred band intensity from (Fig.7.E.C). E). TAP-purified SWI/SNF complex sliding activity using 36W36 Nucleosome in the presence/absence of Runx3 evaluated on. **F).** Quantitative In-Cis remodleing assay using EpiDyne-FRET nucleosomes (15 nM), incubated with SWI/SNF subunits (SMARCA2 /4) as indicated in the presence/ absence of 0.8μM of Runx3 and 1mM ATP. Data are presented as the mean of the Cy3/Cy5 ratio. Quantifications were performed for (N=3).

To address the functional connection between SWI/SNF and Runx3, we investigated whether Runx3 plays any role in SWI/SNF ATP-dependent remodeling activities *in-cis and in-trans*. To do this, we made use of a standard functional remoldeing assay analysing the ability of SWI/SNF to assemble nucleosomes by transferring histone octamers from a doner template (Chicken polynucleosoes) to an acceptor (601-Widom) Cy3 labelled DNA (Fig.7.C). Complete nucleosome assembly indicated by the shift of Cy3-labbeled acceptor DNA template was detected when SWI/SNF subunits (SMARCA2 and SMARCA4) were assisted with Runx3 in the presence of ATP (Fig.7.C, lane 9). Most strikingly, separately SMARCA2 and SMARCA4 were able to partially transfer histone octamers from the donor to the Cy3 DNA template in an ATP-dependent manner only when Runx3 was added to the reaction (Fig.7.C, lanes 5 and 8 consecutively). Quantification of the reaction demonstrated the significant influence of Runx3 in increasing the efficiency of SWI/SNF catalytic activity for nucleosome assembly *in-trans* (Fig.7.D).

Using two additional functional assays, we attempted to analyse another remodeling function of SWI/SNF, which is nucleosomes repositioning or sliding (movement in-cis) that regulates DNA access and consequently gene expression. To do so, we tested the ability of full length SWI/SNF complex (S5.B-C) purified by Tandem Affinity Purification (TAP) in classic sliding assay with cy3-36W36 Widon nucleosomes. Interestingly, SWI/SNF complex displays increased in-cis nucleosomes repositioning activity in the presence of Runx3 as evident by the formation of multiple slow phasic migrating bands (Fig.7.E, lane 3). To further examine this notion, we used EpiDyne-FRET comprised of 5’ Cy3-labelled DNA wrapped around a terminally positioned histone octamer (H2AT120C*Cy5). The activity of an ATP-dependent remodeler (e.g. SWI/SNF ATPase) is detected by a reduction in FRET signal as the Cy3-labeled DNA 5’ end is moved away from the Cy5-labeled octamer. We detected that SMARCA4 ability to catalyse the repositioning of nucleosomes from one location to another was moderately enhanced upon addition of Runx3 (Fig.7. F, red line) compared to the condition with ATP (Fig.7.F, yellow line). Remarkably, the extent of SMARCA4-mediated repositioning was greater during the combinatory effect of Runx3 and ATP (Fig.7.F, green line). This suggested that the Runx3 synergistically enhanced the ability of SMARCA4 to slide nucleosomes across DNA, and this effect was most striking in the presence of the cofactor ATP. We have also observed that Runx3 alone is not capable of catalysing of initiating any nucleosome remodeling activity (Fig.7.F, orange line). These observations indicated that Runx3 activity in SMARCA4-governed chromatin alterations might be in the form of reaction catalysts or a cofactor similar to SWI/SNF classic activators VP-16 or GCN4 (33).

### Runx3 Null Expression Induces Defects in DNA Repair Elicited by Detected Chromatin Abnormalities

Growing evidence has implicated RUNX proteins (Runx1, 2 and 3) as regulators of DNA damage response (DDR), primarily in conjunction with the p53 or Fanconi anemia pathways (35). In addition, DNA damage recognition is associated with chromatin alterations and by changes in post-translational modification pathways such as phosphorylation, methylation and ubiquitination (36). Based on these observations, we tested the hypothesis that Runx3 ablation has effects on cellular functions of the metastatic mesenchymal gastric cancer such as proliferation or impaired DNA damage response secondary to reported chromatin modulations. To this end, the cell proliferation growth curves of HGC-27 cell lines showed moderate higher growth rates in the WT in comparison to Rx3KO (S8.A). The difference was observed after 30hr and become significant reaching 40hr (P<0.05). Next, we assessed response to DNA damage after treatment with 0.5μg/ml Doxorubicin (Doxo). Clear morphological changes such as loss of attachment, irregular cell structure, nuclear defects and cell death were observed in Rx3KO post Doxo treatment (S8.B, lower right panel). We utilized comet assay that detects single and double-strand breaks by measuring leaked DNA tail from single nucleus after electrophoresis, at baseline and after treatment with 0.5μg/ml Doxorubicin (Doxo). This assay gives a comet-like image with the intensity of the tail corresponds with the degree of damage detected in the cells. A low-level halo of DNA damage was observed in both WT and Rx3KO untreated control cells at baseline (Fig.8.A, left panels). Rx3KO cells had significantly higher progressive increase in ‘comet tail’ length indicative of DNA damage, whereas WT cells had significantly smaller ‘tails’ (Fig. 8.B). The average ‘tail DNA length’ after Doxo treatment in Rx3KO cells was 16.33 µm whereas ‘tail length’ in WT cells was only 5.3 µm (Fig.8.C). HGC-27 cells deficient in Runx3 also showed a higher phosphorylation level of γ-H2AX at Ser 139, which is another DNA damage marker required for the assembly of DNA repair at the sites of damaged chromatin (Fig.8.D). Significant increase in γ-H2AX foci was detected in Rx3KO nuclei compared to WT (Fig,8. E), supporting Runx3 potential role in chromatin-dependent genomic integrity. These results indicate impaired DNA damage repair in cells defective of Runx3 expression, and suggesting that RUNX3 might be an integral component of the DNA Damage Response (DDR) through chromatin regulation.

**Fig.8.**
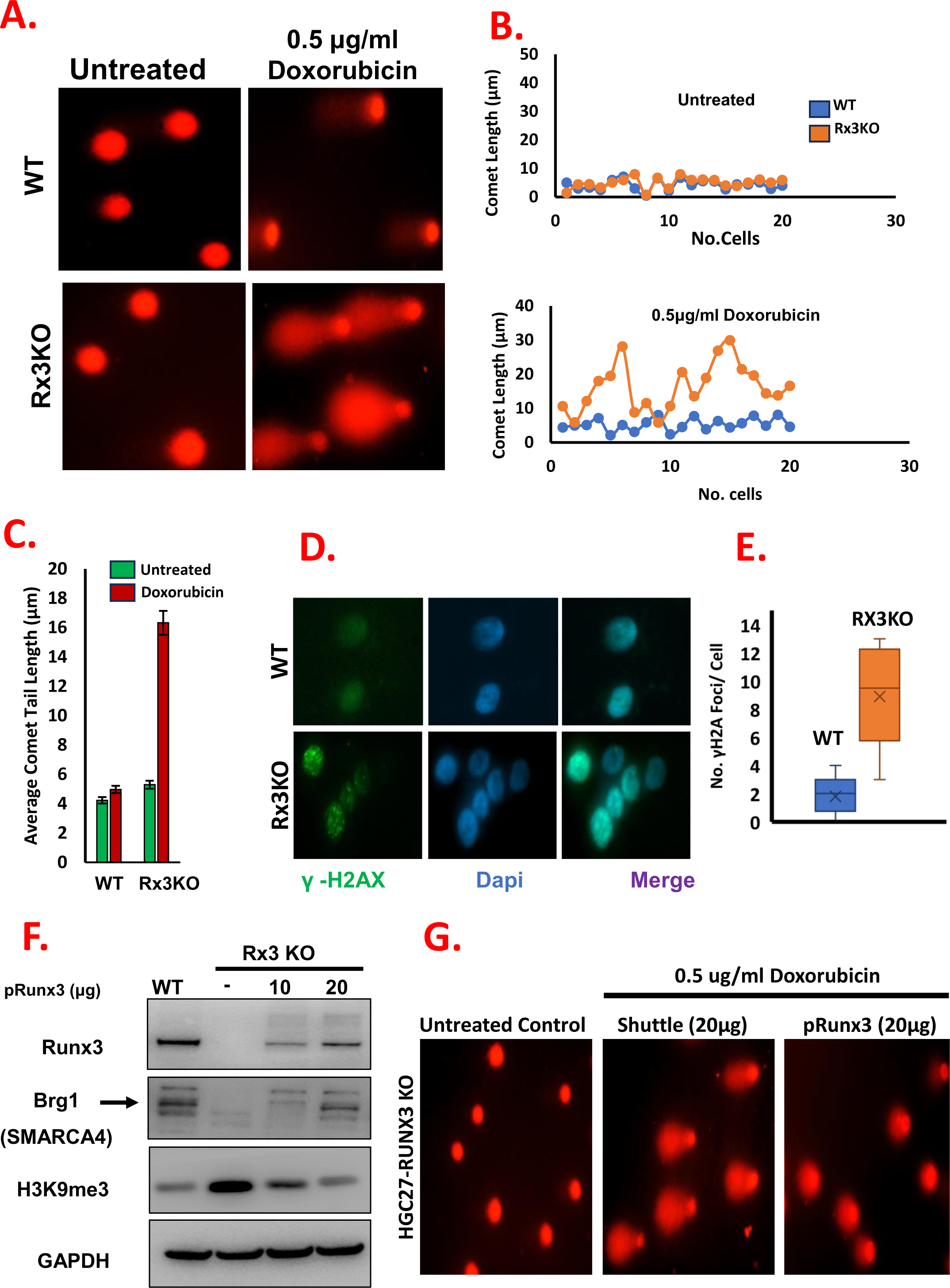
Loss of RUNX3 leads to impairment in DNA damage response rescued by over-expression of Runx3 in Rx3KO. **A).** Increased DNA damage in Rx3KO cells compared with WT cells at baseline and 24hr following 0.5μg/ml Doxorubicin treatment. Images represent individual cell nuclei with tail length directly proportional to damaged DNA degree. Rx3KO cells show progressively more damage compared to WT. **B).** Assessment of ‘comet tail’ lengths for using unified scale for untreated WT and Rx3KO cells (upper panel) and Doxo treated cells (lower panel) showing that Rx3KO cells have significant impairment in their ability to repair DNA whereas WT were able to repair DNA more effectively. **C).** Quantification of average comet tail length. **D).** Induction of γ-H2AX phosphorylation in Rx3KO at baseline compared to WT. **E).** Quantification of the relative no. cells in E, the data shown represent the mean ± SD of three independent experiments. **F).** Immunoblotting of cell lysates from WT and Rx3KO with different levels of Runx3 using anti-Runx3, Anti-Brg1/SMARCA4, anti-H3K9me3 specific antibodies and GAPDH as loading control. These blots are representatives of three independent experiments. **G).** Increased Runx3 re-expression protects against Doxorubicin-induced DNA damage in Rx3KO cells. Forty-eight hours after transfection with transfected with shuttle- or pRunx3-plasmids, cells were treated with 0.5μg/ml Doxorubucin and subjected to DNA damage analysis using comet assay. Quantifications were performed for (N=3).

### Exogenous Expression of Runx3 Restores Chromatin Profile and Recues DNA Repair Response

To further explore the magnitude of Runx3 role in the induced defects in DNA repair elicited by abnormalities in chromatin regulation, we restored adequate levels of Runx3 in Knockout cells by overexpressing CRISPR resistant Runx3 plasmid (S8.C), and measured the impact on some of the reported observations. Increasing Runx3 protein level in Rx3KO cells resulted in a progressive upregulation of Brg1/SMARCAD4 with concomitant decrease in H3K9me3 expression (Fig. 8.F). Next, we tested whether elevated levels of Runx3 can rescue the DNA damage sensitivity. Rx3KO cells transfected with a shuttle vector showed damaged DNA after 0.5μg/ml of Doxorubicin treatment by the presence of ‘tails’ measured by Comet assay (Fig.8.G, mid panel) compared to untreated (Fig.8.G, first panel). On the contrary, Rx3KO cells expressing high levels of Runx3 showed minimal DNA damage after Doxo treatment (Fig.8.G, last panel). Overexpression of Runx3 in knockout cells reversed the DNA damage as indicated by the reduction in the comet length of the majority of transfected cells after Doxo treatment compared to the shuttle group (S8.D). The average ‘tail’ length was ∼13µm in patient cells overexpressing Runx3 compared with 20µm in cells transfected with shuttle vector control (S8.E). Restoring Runx3 expression also resulted in normalization of nucleosomal occupancy profile in Rx3KO cells with increase in mononucleosomes populations associated with reduction in higher polynucleoosmes after MNase digestion for 15min (S8.F). These results indicate that adequate levels of Runx3 are sufficient to regulate proper chromatin architecture and is also required for adequate DNA damage response.

## Discussion

Runt Related Transcription (RUNX) proteins are core regulators of diverse cellular processes and they have been shown to exert paradoxical activities in cancer pathogenesis (37). Despite functional redundancy of the three orthologs, RUNX3 exhibits unique context-dependent biochemical properties and post-translational modifications related to its subcellular localization than RUNX1 and RUNX2 (38). In this study, we demonstrate a novel function of nuclear Runx3 that forms a homodimeric structure directly binding to chromatin in a cooperative fashion. We show that Runx3 modulate chromatin condensation and act as a functional activator enhancing multiple SWI/SNF ATP-dependent chromatin remodeling activities. Additionally, our study shows that Runx3-chromatin alterations are associated with defects in DNA repair machinery.

Here, we hypothesized that the dual functionality of Runx3 in cancer might be location-associated phenomenon. Although the three members of the RUNX family share a high degree of homology, only RUNX3 exhibits cytoplasmic mislocalization “cytoplasmic sequestration” that leads to functional differences and greatly limits its role in gene regulation (39). This in addition to frequent inactivation of RUNX3 in gastric cancers by hemizygous deletion and hypermethylation of its promoter region (8, 40), prompted us to utilize HGC-27 gastric cancer cell line that displays high level of strictly nuclear Runx3 expression as our study model.

All RUNX protein including Runx3 has been reported to heterodimerize with a non-DNA binding core binding factor beta (CBF-β) subunit (41). Our unique observation of Runx3 homodimerization and cooperative binding is similar to those reported in other Snf2 chromatin complexes such as Fun30/SMARCAD1 (42), or Chromatin assembly factor 1 (CAF-1) (43). This indicated that Runx3 might be capable of forming different dimeric/ multimeric structures or levels as in other chromatin remodelling factors (44), or postulated towards its potential to form a prion-like chains. Remarkably, our study aligns with the earlier report that RUNX1 can homodimerizes to assist in folding of distant chromatin sites (45), which suggest that this characteristic might be a common under looked feature of the RUNX family. Targeting of Runx3 to modified nucleosomes, directed us to speculate that it might be recruited by the presence of a specific post-translational modification, yet to be characterized. Although it is intriguing that Runx3 is linked to chromatin binding and nucleosome positioning, further studies are required to reveal the nature of this association.

We can deduce that Runx3 regulation is taking place through two possibilities: 1. Its classic transcription factor role by binding to the promoter or enhancer elements regions, or 2. by chromatin modulation and enhancement of SWI/SNF chromatin remoldeing activities that allow transcription machinery access into regulatory regions. In this study, Runx3 was found to be especially proficient in catalyzing SWI/SNF ATP-dependent remodeling activities. This newly identified function of Runx3 is similar to those observed in the classic activators herpes virus VP16 protein and the yeast Gcn4 shown to target SWI/SNF and enhance its mechanical chromatin mediated transcriptional regulation (46). Several reports indicated that human SWI/SNF complex can be recruited to target genes through direct interactions with these sequence-specific transcription activators such as c-MYC and contribute to cancer aggressiveness (47). Unlike the targeting by Myc governed by transcriptional regulation, our data indicate that Runx3 shares functional analogy with VP16 and GCN4 in acting as cofactor stimulating SWI/SNF structural chromatin changes such as octamer transfer and sliding activities.

Interestingly, we detected a functional and causal link between Runx3 and the catalytic subunit SMARCA4 of SWI/SNF complex. The proteolytic-ubiquitin dissolution of SWI/SNF subunits observed in Runx3 null expression cells, supported the reported role of Runx3 in facilitating ubiquitination and proteasomal degradation of several proteins (48). Additionally, the physical interaction of the two proteins along with the co-expression correlation observed in SW13 cell line and patient’s cohort analysis indicated a potential regulatory feedback loop between Runx3 and SWI/SNF associated with cancer stage and degree.

Runx3 appears to be special in that it affects silencing through increased chromatin condensation and HP1 oligomerization, which strengthen its novelty as a potential heterochromatin regulator possibly as a boundary element such as Fun30/SMARCAD1(49–50). Our work proposes that Runx3 regulation of each HP1 isoform makes a distinctive contribution to the organization and structure of heterochromatic foci. These individual signatures mediated by the loss of Runx3 in HGC-27 support the previous observations on the role of HP1 isoforms and their functional link with heterochromatin structure and genome organization despite a well-established functional redundancy between despite a well-established functional redundancy between these isoforms (29). We reasoned the reduction in HP1α level due to RUNX3 ablation would reveal its relative contributions to cancer oncogenicity as the level of HP1α overexpression significantly correlates with global survival and the occurrence of metastasis (30).

We attempted to obtain further insight into the biological function of Runx3-mediated chromatin alterations by identification of dysregulated pathways and interaction partners. Our proteomics analysis revealed that Runt domain of Runx3 is crucial for maintain interactions with multiple chromatin organization and SWI/SNF subunits. This supported the previous studies of runt domain mediated transcriptional regulation and physical interactions with the bromodomain of BRD2, and SWI/SNF protein complexes to promote chromatin opening (51). Through integration of ATAC-seq and RNA-seq, several inflammatory response pathways were upregulated while Myc targets, proliferation and DNA repair pathways were downregulated. These data supported studies of RUNX3 targeting MYC protein and their dynamic functional association (52). Analysing the ratio between potential RUNX3/MYC and the reported MYC/MAX or MYC/MIZ-1 interactions in this context will uncover additional insights to better understand how Runx3 can exhibit both oncogenic and tumor-suppressive properties.

On the other hand, and since HGC-27 cells are defective for WT p53 activity, we can propose that p53 deficiency causes RUNX3 to become an oncogene, resulting in abnormal upregulation of MYC (53). This notion is validated by the reduced Myc expression in Rx3KO cells using integrated RNA-seq analysis (Fig. 5.B, red section), and immunoblotting analysis of c-Myc (Fig.S2.D). It is interesting to note here that mutant p53 forms were shown to result in gain of functions in the form of upregulated chromatin pathways driving cancer growth (54). This suggest that Runx3 mediated chromatin regulation might be due to alterations of TP53 in HGC-27. This highlights the critical role of the Runx3 in maintenance of genomic integrity possibly through the highly interconnected relationship between RUNX factors and the p53 pathway (35). Our results show reduced repair of DNA lesions in Runx3 knockout HGC-27 cells associated with increase in phosphorylation of γ-H2AX. Our work provides new view of Runx3 involvement in maintaining genomic stability and responding to cellular stress via chromatin mediated regulation. This newly described function of Runx3 as a chromatin factor indicate that it might be the trigger of its oncogenicity and its potential to be a multifunctional protein.

This study together shows clear cellular defects in DNA damage repair in Rx3KO cells with aberrant chromatin architecture that might impede DNA repair machinery from accessing the damaged lesions. Although this strengthen the link between impaired Runx3-mediated DNA damage response and chromatin, it appears that this connection is more complex than previously thought and additional work is needed to decipher and elucidate this interesting direction. Our results point toward robust physical association between Runx3 and multiple Snf2-related chromatin factors including SWI/SNF. Thus, it will be important to evaluate in the future the detailed kinetics of these interactions and their involvement in the Runx3 mediated gene regulation. Overall, the data we present in this study reveal that Runx3 is a new homodimeric chromatin modulator in addition to its previously reported transcription factor activities, supporting the conceptualization of its dual functionality as both a tumor suppressor and oncogene. Runx3-chromatin axis is vital for homeostasis of several molecular mechanisms including nucleosome positioning, heterochromatin maintenance and DNA damage response. Which imply a promising direction to screen for anti-Runx3 epigenetic drugs. However, it remains to be seen whether and how these processes also converge in the molecular pathogenesis of Runx3 dependent cancers.

## Materials and Methods

### Cell culture

HGC-27, IM95, AGS, MK45 and MKN28 were purchased from CellBank Australia. SW13 was acquired from American Type Culture Collection. While HFE145 the immortalised human normal gastric epithelial cell line HFE-145 was a gift from Drs. Hassan Ashktorab and Duane T. Smoot (Howard University, Washington, DC, USA). All cell lines were cultured in RPMI-1640 Medium (Nacalai tesque), supplemented with 10% fetal bovine serum (FBS) (Biowest) and 1% Penicillin–Streptomycin (Gibco by Life Technologies), except SW13 which was maintained in 10% DMEM media. Cell lines were authenticated by DNA profiling (Promega). All cell lines were tested regularly to be mycoplasma-free by Mycoplasma detection assay (Sigma Aldrich).

### Genome engineering by CRISPR/Cas9

Two biological knockout clones were created from the sgRNA sequence including PAM region targeting exon 3 of RUNX3 which was determined by the CRISPR Design Tool (http://www.genscript.com/CRISPR-gRNA-constructs.html) as follows: 5’-GGACGTGCCGGATGGTACGG TGG-3’. Gene-specific sgRNA oligos were cloned into a pLenti CRISPR v2 plasmid (Genscript), which bicistronically expresses Cas9 nuclease. HEK293T cells were co-cultured with 15 μL of TransIT-LT1 (Mirus Bio) reagent and 5 μg of plasmids comprising sgRNA and packaging vectors PLP1, PLP2, and PLP/VSVG (Addgene). After 48 - 72 hours, virus-containing supernatant was added to the target cells in the presence of 5µg/mL polybrene (Sigma-Aldrich). After 72 hours, Puromycin (Invitrogen) was added for 5 days to select for transduced cells. RUNX3 deletion was confirmed at mRNA and protein levels after single cell cloning, and two distinct clones (K1 and K2) with complete loss of Runx3 were selected for this study. Rx3KO was the designated term used to identify the knockout from Wild Type (WT).

### Biotin pulldowns and Proteomics

Using the TurboID methods (Branon et al., 2018), proteins proximal to RUNX3 and TurboID (control) were biotinylated and isolated using streptavidin-bead pulldowns. pRetroX-3xHA-T*urboID-RUNX3-Puro or* pRetroX-3xHA-T*urboID-RUNX3(R122c)-Puro* or pRetroX-3xHA-T*urboID-*Puro were transduced in TetOn-HGC-27 cells and a stable population was selected. Briefly, 24 hr after induction with 1µg/mL Doxycycline, medium was supplemented with biotin at 50μM. Cells were collected 48 hr after transfection, washed 3x on ice with cold Phosphate Buffered Saline (PBS) and scraped in lysis buffer (LB), prepared in RIPA buffer (Thermo): 1x Protease inhibitor cocktail (Roche), 1x Halt™ Phosphatase Inhibitor Cocktail (Thermo), 1x Benzonase (Novagen), 1mM PMSF, 10µM MG132; 1ml per 10-cm plate]. Cell lysates were incubated overnight with 60μl of equilibrated Dynabeads MyOne Streptavidin C1 (Invitrogen). Beads were subjected to stringent wash using Wash buffer (WB), prepared in PBS: 150mM NaCl, 10mM Tris (pH 7.4), 1% Triton-X, 1mM EDTA, 0.2mM Sodium orthovanadate and 1x Protease inhibitor cocktail (Roche). For elution of biotinylated proteins, beads were heated at 98°C in 50μl of Elution Buffer (EB), prepared in lysis buffer: 1x Laemmli buffer and 1mM DTT. Beads were separated using DYNAL Bead separator rack (Invitrogen). For mass spectrometry, 100 μg of HGC-27 At and Rx3KO were lysed in RIPA buffer on ice for 10 min following published protocols (55).

### Expression and Purification of Recombinant Proteins

In house recombinant Runx3 protein was produced from His-tagged RUNX3 plasmid. Briefly, RUNX3 was amplified from human genomic DNA and subsequently subcloned into the pET21D expression plasmid. pET21D was then engineered to contain an in-frame hexa-histidine tag (SSHHHHHH; 6xHis) at the C-terminus of the RUNX3 gene (pET21D6H). His-tagged RUNX3(C-terminal 6XHis) was expressed from pET21D6H in the Rosetta2 E. coli strain (Novagen) at 37 °C until OD600 reached log phase 0.5, then cells were harvested by centrifugation and resuspended in lysis buffer (20 mM Tris pH 7.5, 350 mM NaCl, 0.05 % β-mercaptoethanol) containing protease inhibitors (0.2 mM AEBSF, 2 µM E64, 2.6 mM aprotinin, 1 µM pepstatin) with 0.1 % Tween-20. Cells were lysed by freeze/thawing in liquid nitrogen and sonication. The soluble fraction was extracted by centrifugation at 35000 g at 4°C for 30 min. Runx3-6xHis was then purified in batch over HisPur Ni-NTA Magnetic beads. The Runx3-6XHis was eluted using elution buffer (100mM sodium phosphate, 600mM sodium chloride, 250mM imidazole). Runx3-6XHis was then concentrated in 50 KDa MWCO centrifugal concentrators and dialyzed against 20 mM Tris pH 7.5, 350 mM NaCl, 10% Glycerol, 1 mM DTT and stored at -80 C. Recombinant Runx1(ab134873) and Runx3 (ab95893) and CBFβ (Abcam: ab98252) for comparison were purchased from Abcam.

### Fluorescent Labelled Mononucleosomes Assembly

Nucleosomes were assembled by mixing equimolar amounts of histone octamer (Epicypher) modified or recombinant as indicated and DNA in high salt and performing stepwise dialysis into low salt as described (56). DNA was generated by preparative PCR using primers obtained from IDT (Singapore) fluorescently labelled where appropriate. Nucleosomes were assembled on DNA fragments based on the synthetically selected Widom-601 sequence (26). We have adopted a nomenclature in which the lengths of DNA extensions on either side of a nucleosome are indicated as numbers on either side of a letter that defines the core positioning sequence used. So 36W36 designates a nucleosome positioned on the Widom-601 sequence with a 36 bp extension on both (upstream/ down-stream) sides of the DNA fragments. The oligos used to generate the 36W36 fragment5′GGCGAATTCGAGCTCGGTAC and 5′AGGTCGACTCTAGAGAATCC. The PCR fragments were purified by PCR purification kit (QIAGEN).

### Quantitative Electrophoretic Mobility Shift Assay

Recombinant His-Runx3 and occasionally Runx1 (Abcam: ab134873) or CBFβ (Abcam: ab98252) were serially diluted in 1X HMA buffer (20 mM HEPESpH 7.6, 25 mM KOAc, 5 mM MgAc) containing 0.1% Tween-20. Binding reactions were established in a final volume of 20 µL containing 0.5 X HMA buffer, 0.1 mg/ml BSA, 1 mM DTT, 5 mM AEBSF, and 30 nM Cy3 labeled nucleosome or DNA and 2 µL of Fun30-6xHis diluted to the concentrations described in the figures. Reactions were allowed to equilibrate on ice for 30 min before electrophoresis on 0.5X TBE 6.5% native polyacrylamide (37.5:1) gels at 130 V for 50 min at room temperature. Gels were scanned on Typhoon Trio laser scanner (Amersham Bioscience) for the Cy3 signal and the nucleosome/DNA bands were quantified ImageJ.

### Octamer Transfer and Nucleosome Sliding Assays

The same template 36W36 used in binding reactions, was utilized in these assays. For octamer-transfer assay, 30 ng of Cy3-naked DNA template was used as an octamer acceptor in this assay, was incubated with ∼10 ng of donor nucleosomes (Epicypher) in the presence of the indicated proteins (Runx3, SMARCA2, SMARCA4) and ATP for 2 h at 30 °C in the same binding buffer as before. As for the sliding assay remodeling reactions contained 30ng of Cy3-W36W assembled nucleosomes, 1 mm ATP, 3 mm MgCl2, 50 mm NaCl, 50 mm Tris-Cl, pH 8.0, TAP-SWI/SNF and Runx3 as indicated. Reactions were incubated at 30 °C for 30 min and terminated by addition of 0.5 μg of λHindIII-digested DNA and quenching on ice. Sucrose was added to a final concentration of 2% as a loading buffer for both assays and reactions were run and resolved on 6.5% Native Poly Acrylamide Gels (49:1) acrylamide to bisacrylamide ratio at 150 volts for 1.5 h. The gels were visualized by Typhoon Trio laser scanner (Amersham Bioscience).

### FRET Nucleosome Remodeling Assay

To measure real-time nucleosome remodeling induced by enzymatically active remodeling complexes SMARCA4, SMARCA2, and our candidate factor Runx3 we utilized The EpiDyne®-FRET mononucleosome (Epicypher) following manufacturer instructions. Briefly,E piDyne-FRET nucleosomes (20 nM) were incubated with 10 nM of indicated proteins (SMARCA2/4, Runx3) and combinations in the presence or absence of 2 mM ATP. Upon ATP addition, reactions were read in TECAN-SPARK plate reader at times indicated. Data is presented as the mean±SD of the Cy3/Cy5 ratio (N=6).

### MNase Digestion

Cell pellets were first resuspended in permealisation solution-I with triton X-100 (150 mM sucrose, 80 mM KCl, 35 mM HEPES pH 7.4, 5 mM MgCl2, 0.5 mM CaCl2, 0.05% triton X-100) at room temperature for 3 mins. It was followed by centrifuging and washing with perabilisation solution-I without triton X-100. The cells were then resuspended in permeabilisatoin solution-II (150 mM sucrose, 50 mM Tris-Cl pH 7.5, 50 mM NaCl, 2 mM CaCl2) containing micrococcal nuclease (Theromo Scientific) and incubated at room temperature for 5 mins. The enzymatic reaction was stopped using the equal volume of stop solution (20 mM Tris Cl Ph 7.4, 0.2 M NaCl, 2 mM EDTA, 2% SDS, 0.2 mg/ml proteinase K) and incubate at 37C water bath for 2hrs. The DNA extraction was performed by adding equal volume of phenol-chloroform isoamyl alcohol and it was centrifuged at 13000rpm for 30mins at room temperature. DNA was then precipitated by adding 2X volume of isopropyl alcohol and centrifuged at 12000 rpm at 4C for 15 mins. The supernatant was removed and the white pellet was resuspended in 40ul of TE buffer. The DNA was checked on 3% agarose gel at 50V for 90mins or when there is a good separation.

### Western Blotting

Whole cell extract was obtained by lysing cell pellets using RIPA buffer followed by gentle sonication using Water bath Bioraptor to release chromatin bound proteins. The concentration of protein lysates was measured by Bradford assay using the BioRad protein detection kit (Bio-Rad). Typically, 30µg of the whole cell extract was analyzed in Western blots. Western blot analysis was performed by running a 4-12 % NuPAGE® Novex® Bis-Tris Gels (Invitogen, USA), then transfer of proteins to a PVDF membrane at 200 V for 1 hour in NuPAGE® 1X transfer buffer using a novex transfer chamber. The membrane was then blocked in 50 ml of TBS–Tween (50 mM Tris, 138 mM NaCl, 2.7 mM KCl, 0.05% Tween 20, pH 8.0) containing 5 % milk at 4 °C overnight. The membrane was then incubated with 1:1000 dilution of primary antibodies as listed in the blot figures in PBS for 2 hrs at room temperature. The membrane was then washed three times for 10 min each in of TBS–Tween, and incubated for 1 h with 1:3000 of the corresponding secondary antibodies.

### Immunofluorescence Microscopy

Cells were plated on coverslips in six-well dishes at a density of 0.5 × 10^6^ cells/well. After fixation in formaldehyde, cells were permeabilized and stained with indicated antibodies followed by fluorescein isothiocyanate-conjugated anti-rabbit immunoglobulin G (Sigma, dilution 1:100). The cover slides were mounted with Vectashield mounting media containing 4′,6-diamidino-2-phenylindole (Vector Laboratories). Signals were observed by fluorescence microscopy using Zeiss microscope.

### Migration and Wound Healing Assays

To measure cell migration activity, Polystyrene plates with 6.5mm Transwell with 8.0µm pore polycarbonate membrane insert (Corning) were used. For invasion assay, the surface of upper chamber was additionally coated with 50 µL of Matrigel (Corning). 4 × 105 of cells suspended in 200 μL of RPMI-1640 with 0.1% BSA were added to the upper chamber. Lower chambers were filled with medium containing 20% FBS as chemoattractant. After incubation at 37°C for 12 - 48 hours (according to cell type), the cells that migrated / invaded to the lower side of the upper chamber were counted. Non-migrated cells were removed by swabbing top surface of the membrane insert. Membrane containing invading cells was fixed with methanol, and stained with hematoxylin (3 minutes) and eosin (1 minute), and mounted on slides. The migrated cells were counted under light microscope for 5 fields randomly. For wound healing HGC-27 cells were seeded in 24-well plates at 2 × 10^5^ cells/mL and incubated for 24–48 h. After cells reached 100% confluence, wounds were generated using a 1 mL micropipette tip. Media was removed, cells washed with 500 μL PBS, and 500 μL of complete culture media containing compounds added into each well. Images were acquired immediately following media replacement (T = 0), and after 6, 12 and 24hr at 10×. After exporting images, wound areas were measured using ImageJ. Briefly, a polygon selection tool was used to indicate wound area, and quantified (*Analysis > Measure*). Extents of closure at T6, T12, T18, and T24 were calculated by subtracting area at T0; percentage closure was determined by normalizing difference to area at T0.

### Proteasome inhibitor treatment

To examine proteosomal degradation, cells were treated with dimethylsulfoxide (DMSO) or 40 µM MG-132 proteasome inhibitor for 12 and 24 h. Cell lysates were analyzed by western blotting with the corresponding antibodies as described before.

### RT 2 Profiler PCR Arrays

Total RNA was extracted from WT and Rx3KO HGC-27 cells and analyzed by Qiagen R2 Profiler kits: Human chromatin remoldeing factors arrays (PAHS-086Z) and Human chromatin modification enzymes arrays (PAHS-085Z) following manufacturer instructions.

### Comet Assay

The comet assay (alkali method) was modified from that described earlier (57). All of the steps (preparation of slides, lysis and electrophoresis) were conducted under red light or without direct light in order to prevent additional DNA damage. After electrophoresis, the slides were placed in a horizontal position and washed three times (5 min each) with 0.4 M Tris buffer, pH 7.5, to neutralize the excess alkali. Finally, 70 ml of ethidium bromide (2 μg/ml) was added to each slide, which was then covered with a cover slip, stored in a humidified box at 4°C and analyzed using a Nikon Eclipse Ti-E fluorescence microscope under ×40 objective with a calibrated ×10 eyepiece. Images of 50 randomly selected cells were analyzed from each sample. Tail lengths (nuclear region + tail) were measured at ×40 magnification in which one pixel unit was ∼0.085 μm. The fluorescence microscope was equipped with a BP546/12-nm excitation filter and a 590-nm barrier filter. Cells were also scored visually into two classes, according to tail size (from undamaged to damaged). The final overall rating for the DNA damage score was represented as a value of the average of comet length in 50 cells and error bars represents standard deviation.

### Immunoprecipitation (IP) and Co-IP

Total cell protein extracts were incubated with Dynal protein A magnetic beads (Invitrogen) coupled to the indicated antibodies in a pulldown buffer (50 mM N-2-hydroxyethylpiperazine-N′-2-ethanesulfonic acid [pH 7.5], 1 mM EDTA, 150 mM NaCl, 10% glycerol, 0.1% Tween 20, 0.5 mM dithiothreitol (DTT), 1 mM phenylmethylsulfonyl fluoride, 2 μg/ml leupeptin and 2 μg/ml pepstatin A). Cell extracts were incubated with the beads overnight at 4°C while mixing on a rotating wheel. Beads not conjugated with antibody were used as negative control. Supernatants were collected, and the beads were then washed with pulldown buffer and left as a 50% slurry. Bead fractions were analysed on 4–12% SDS-PAGE followed by immunoblotting detection using the corresponding antibodies. Coimmunoprecipitation from a the Rosetta2 E. coli strain expressing either His-Runx3, GST-Runx3 using pGEX4T-1-RUNX3 aa 49-187 (Addgene) or both His- and GST-tagged Runx3. Followed by immunoprecipitation using the corresponding antibodies and western blotting analysis of bound vs supernatant (unbound) fractions on 4-12% SDS-PAGE.

### Over-expression of RUNX3

*RUNX3* expression vector containing CRISPR resistant sequence was designed using Vector Builder (ID: VB220609-1019qeh) and produced by Yunchu Bioscience, China. HGC-27 RX3KO cells were transfected with empty shuttle-GFP vector or RUNX3 expression vector using lipofectamine 2000. Twenty-four hours after transfection, cells were washed and maintained in growing media for 48 h. Subsequently, cells were processed for western analysis, comet assay or MNase analysis as described.

### ATAC-Seq

For ATAC-seq, HGC-27 WT and Rx3KO cells were plated at 100,000 cells/ ml density and subjected to downstream hyperactive Tn5 transposase treatment, followed by tagmentation-based NGS library preparation and sequencing was conducting by Novogene Services, Singapore. ATAC-seq data analysis were processed with the nf-core atacseq pipeline (version 2.0). Unless specified, default parameter settings were used. In brief, after adapter removal and read quality trimming with Cutadapt and TrimGalore, 150 bp paired-end ATAC-seq reads were aligned to the GRCh38 genome via the bwa aligner. Duplicate reads were then marked with picard. Reads were filtered according to a list of criteria: whether they mapped to the mitochondrial DNA or the ENCODE blacklisted regions, whether they were marked as duplicates, whether they were soft-clipped et cetera. ATAC-Seq peak regions of each sample were called using MACS2 with the broad peaks setting, and the effective genome size was set via –macs_gsize = 2897310462.20. ChIPseeker was used for plotting peak profiles and for annotating peaks relative to genomic features, with the TSSRegion set to be +-3kb from the TSS. For assessment of peak quality, a custom function was written to compute the TSS Enrichment Score according to the ENCODE project standard. The deeptools package was used to generate the fingerprint profiles. An ATAC-seq specific QC html report was generated with ataqv, and the preceding QC results were collated into a single reportwith multiqc. Peaks that were shared across both peaksets in each biological condition were included to form an overall consensus peakset. Differential peaks were identified using DiffBind with two biological replicates per condition. Read count information for the consensus peaks were obtained from the bam files and analysed using the DESEQ2 method. Following the authors’ recommendation, we set background=TRUE and normalize=DBA_NORM_NATIVE for the dba.normalize function. The fold-changes of peaks were evaluated, and peaks with a FDR <= 0.05 were considered differentially accessible. Profile plots over the differentially accessible peaks were also generated from the DiffBind package. Peaks that were gained or loss from the RUNX3 KO vs WT comparison were the subjected to GREAT for pathway analysis. In our analysis, we created a consensus peak set from peaks that were present in all 5 biological replicates for each condition. we computed the transcription start site (TSS) enrichment score. The TSS score is a ratio between the aggregate distribution of reads centered on TSSs and that flanking the corresponding TSSs. TSS score = the depth of TSS (each 100bp window within 1000 bp each side) / the depth of end flanks (100bp each end). TSSE score = max(mean(TSS score in each window)). TSS enrichment score is calculated according to the ENCODE project standards, TSS enrichment values < 5 are “concerning”, 5-7 are “acceptable”, and >7 are “ideal”.

### ATAC-seq Correlation with The Cancer Genome Atlas Stomach Adenocarcinoma (TCGA-STAD)

We downloaded the TCGA-STAD specific peak sets from the GDC ATAC-seq database. Specifically, a total of 121,582 peaks were obtained from the provided STAD_peakCalls.txt file. To determine the chromosome accessibility score, we grouped the consensus ATAC-seq peaks by chromosome and tallied the number of peaks per chromosome. Following this, we normalised the peak counts by chromosome length and to ensure that scores ranged between 0 and 1, we further scaled these normalised counts by their cumulative sum and a constant scaling factor.

### Integrative analysis of ATACseq and RNAseq data

To determine whether changes in open chromatin regions in ATAC-seq analysis were associated with changes in gene expression, we integrated our ATAC-seq data with RNA-seq data. Total RNA was extracted from WT and Rx3KO cells using Qiagen RNeasy kit following manufacturer instructions. Then RNA samples were analyzed and sequenced by Novogene Services, Singapore. The nf-core/rnaseq (v3.12.0) pipeline from the nf-core collection of workflows, was used to analyse the RNAseq data. In brief, quality control of reads was carried out with FastQC, followed by adapter and quality trimming with TrimGalore. Reads were then aligned to the GRCh38 human reference genome with STAR and transcript quantification was conducted with Salmon. Differential gene expression analysis was performed following the edgeR, Limma and Voom workflow (58). DEGs were identified with a predetermined adjusted p-value cutoff of 0.05. Gene set testing was carried out with the Camera method (59), against the MsigDB hallmark geneset. Enrichment analysis was conducted against the ReactomePA pathways and the MSigDB Hallmark gene set. Prior to DEG analysis, we plotted a multi-dimensional scaling (MDS) plot and discarded a Rx3KO sample as it appears distinct from the other 2 biological replicates. As a result, a total of 2 and 3 biological replicates were used for the Rx3KO and WT condition respectively.

### Transmission Electron Microscopy (TEM)

HGC-27 WT and Rx3KO cells were plated at 50,000 cells/ml and cultured on 12mm slides in 6-well plates until they reach 70% confluency. Samples were then fixed with a solution containing 2.5% glutaraldehyde containing in 0.1M phosphate saline buffer (PBS, pH 7.2), then washed three times in 0.1M PBS and post-fixed for 1h at RT with a mixture of potassium ferrocyanide-osmium tetroxide solution. Samples were dehydrated in a graded series of ethanol and then infiltrated and embedded in Araldite medium. Ultrathin sections were cut using a Leica Ultracut microtome, stained with lead citrate and observed under a JEOL JEM 1400 transmission electron microscopy operated at 100 kV. Digital acquisition was performed with a Matataki Flash sCMOS camera.

### Cyto/Nuc Fractionation

HGC-27 WT and Rx3KO cells were plated at 400,000 cells/ml density and harvested after cells reaching 70% confluency. Cytoplasmic and nuclear fractions were isolated from cell pellets using Abcam Mitochondria/ Cytosol Fractionation Kit (Ab65320) as manufacturer instructions. Then purified fractions were analysed using Western blotting against illustrated antibodies.

### Kaplan-Meier Survival Curves

Kaplan-Meier plotter containing survival statistics over 876 of the gastric cancer patients subjected to mRNA expression profiling, were used to generate survival curves of different probes Runx3, SMARCA4 and their combined expression. (https://kmplot.com/analysis/index.php?p=service&cancer=gastric)

### Spheroid Culture

The detailed protocol is described in (60). In brief, HGC-27 WT and Rx3KO cells (2.5 × 104) were plated on a 1.5% agar bed (w/v) for 3 days. Three-dimensional spheroids were transferred and embedded into collagen I solution. The next day, collagen-embedded 3D tumor spheroids.

Tumor spheroids were extracted from collagen solution using collagenase type 1 (Sigma-Aldrich) for downsteam processing. Pictures were obtained using a Nikon A1R confocal microscope.

### TMA and Immuhistochemistry

Gastric Cancer Tissue MicroArray slide (NBP2-30308) of 59 specimens were purchased from Novus Biologicals and subjected to immunohistrochemistry staining using specific Runx3 antobody ((Santa Cruz, sc-376591, dilution 1:500) based on a published protocol (61). Briefly, traditional DAB immunohistochemical staining was used using the Leica Biosystems Bond Polymer Refine Detection Kit (DS9800). The slides were baked and dewaxed followed by heat induced epitope retrieval (HIER) at 100 °C in antigen retrieval buffer for 20 min. The slides were then peroxidase blocked (only for the 1st marker) for 10 min. Markers were prepared in DAKO antibody diluent followed by the polymeric HRP-conjugated secondary antibody (DS9800) and opal fluorophore-conjugated TSA (Akoya Bioscience) at 1:100 dilution before dispensing to the slide sequentially. Slides were rinsed with 1 × washing buffer after each step.

### Deep Complex and Haddock Protein Docking Analysis

For protein docking and dimerization models, the crystal structures of the proteins were downloaded as PDB format from AlphaFold Protein Structure Database (https://alphafold.ebi.ac.uk/). The UniProt ID was as indicated: For Runx3 (Q13761), while SMARCA4 (Q8R0K1). Then two web tools modelling platforms were used: 1. Deepcomplex (http://tulip.rnet.missouri.edu/deepcomplex/web_index.html) to generate the homodimerization model of Runx3 as per the published study (62), and 2. Haddock 2.4 (https://wenmr.science.uu.nl/haddock2.4/) to predict Runx3 and SMARCA4 heterodimerization (63–64). The predicted molecular complex was the top of 10 clusters conformations with lowest docking energy structure.

### Tandem Affinity Purification (TAP) of SWI/SNF

The wild-type SWI/SNF TAP tag strain was purchased from Euroscarf and protein complex was purified from yeast whole cell extract by tandem affinity purification (TAP) over two affinity columns as described previously (65). Briefly, whole cell extracts were isolated from six liters of yeast cultures grown in YPD media and added to IgG Sepharose 6fast flow resins (Cytiva). The complexes were eluted from the beads by TEV Protease (New England Biolabs/ NEB) cleavage in a buffer containing (10 mm Tris (pH 8), 150 mm NaCl, 0.5 m EDTA, 0.1% Nonidet P-40, 10% (v/v) glycerol, 1 mm phenylmethylsulfonyl fluoride, 2 μg/ml leupeptin, 1 μg/ml pepstatin A, and 1 mm dithiothreitol. Following binding to calmodulin spharose 4B resins (Cytiva), the complexes were eluted using a buffer containing (10 mm Tris (pH 8), 150 mm NaCl, 1 mm MgAc, 1 mm imidazole, 2 mm EGTA, 0.1% Nonidet P-40, 10% (v/v) glycerol, 1 mm phenylmethylsulfonyl fluoride, 2 μg/ml leupeptin, 1 μg/ml pepstatin A, and 0.5 mm dithiothreitol). Purification quality was monitored by Western blot using Snf5 as well as SYPRO Ruby staining (Invitrogen).

### Statistical Analysis

Data are presented as mean ± standard deviation (SD) or standard error of the mean (SEM). Differences between groups were calculated by unpaired two-tailed Student’s t-test; p < 0.05 was considered significant. Calculation of cooperativity of binding curve was measured using Hill Plot.

## Acknowledgment

We thank Dr. Vivien Koh for her assistance in generating the spheroid and 3D culture and all members of Prof. Ito lab for their support and resources sharing. European e-Infrastructure projects are acknowledged for the use of their web portals and utilization of Haddock platform. Also we thank Dr. Kane Toh from Genomics and Data Analytics Core (GeDaC), Cancer Science Institute, Yong Loo Lin School of Medicine, National University of Singapore for providing bioinformatics support.

---

**S1.**
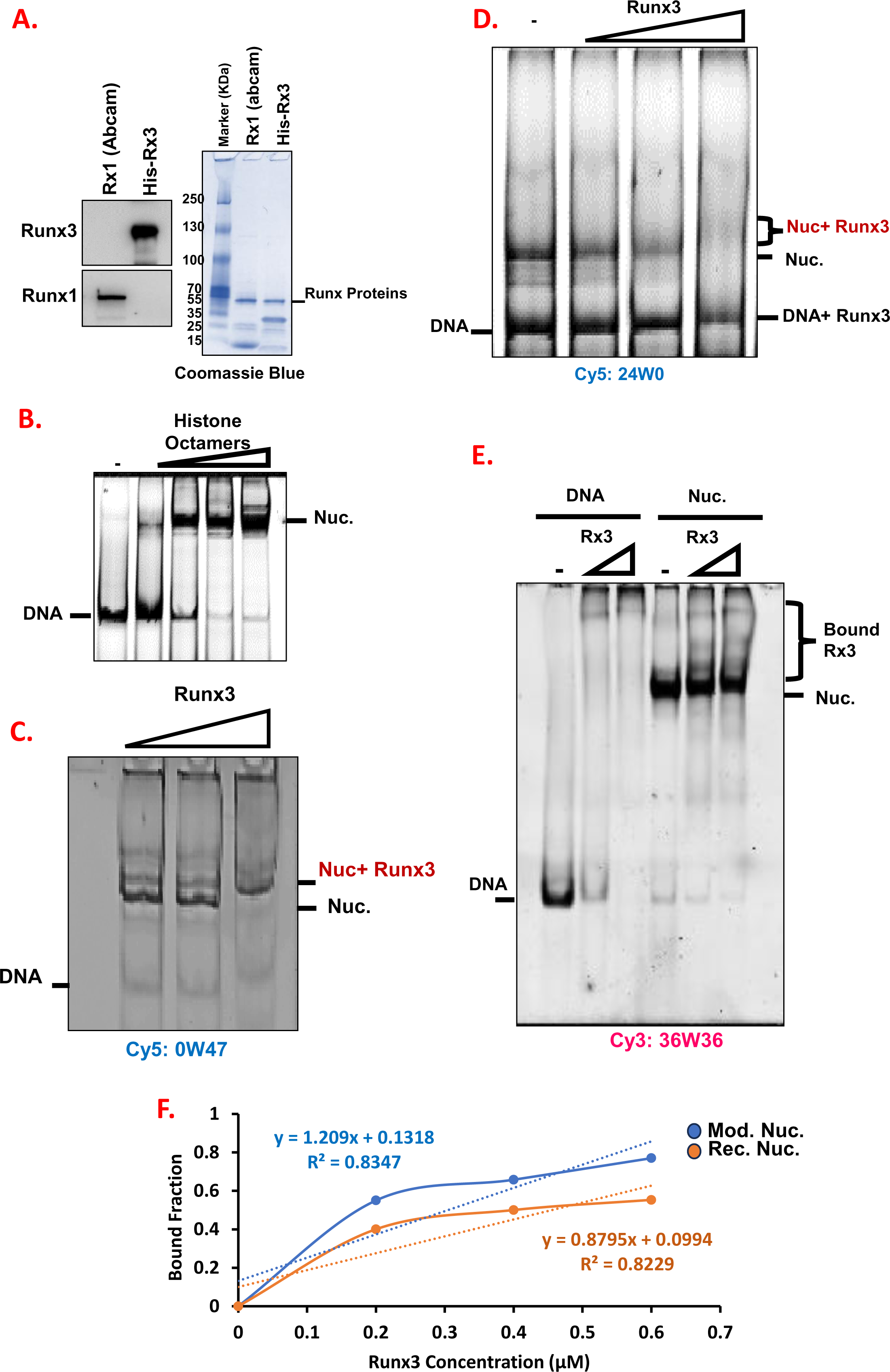
Runx3 binding to different nucleosomes structures and comparisons of affinity mangtitude. **A).** Visualization of purified recombinant His-Runx3 and Runx1 (Abcam) by Commassie staining and Western Blotting using specific antibodies. **B).** Reconstitutions of fluorescent labelled mononucleoosmes at 1 μM concentration by stepwise dialysis. Comparisons of Runx3 binding affinity with: **C).** 0W47 mononucleosomes, **D).** 24W0 mononucleosomes. **E).** Binding of titrated equimolar amounts of Runx3 to DNA and 36W36 mononucleoosmes. In all reactions 30 nm of fluorescent labelled mononucleosomes were used. **F).** Quantification of Runx3 binding affinity to Cy-36W36 assembled into either modified nucleosomes (blue) or recombinant nucleosomes (orange).

**S2.**
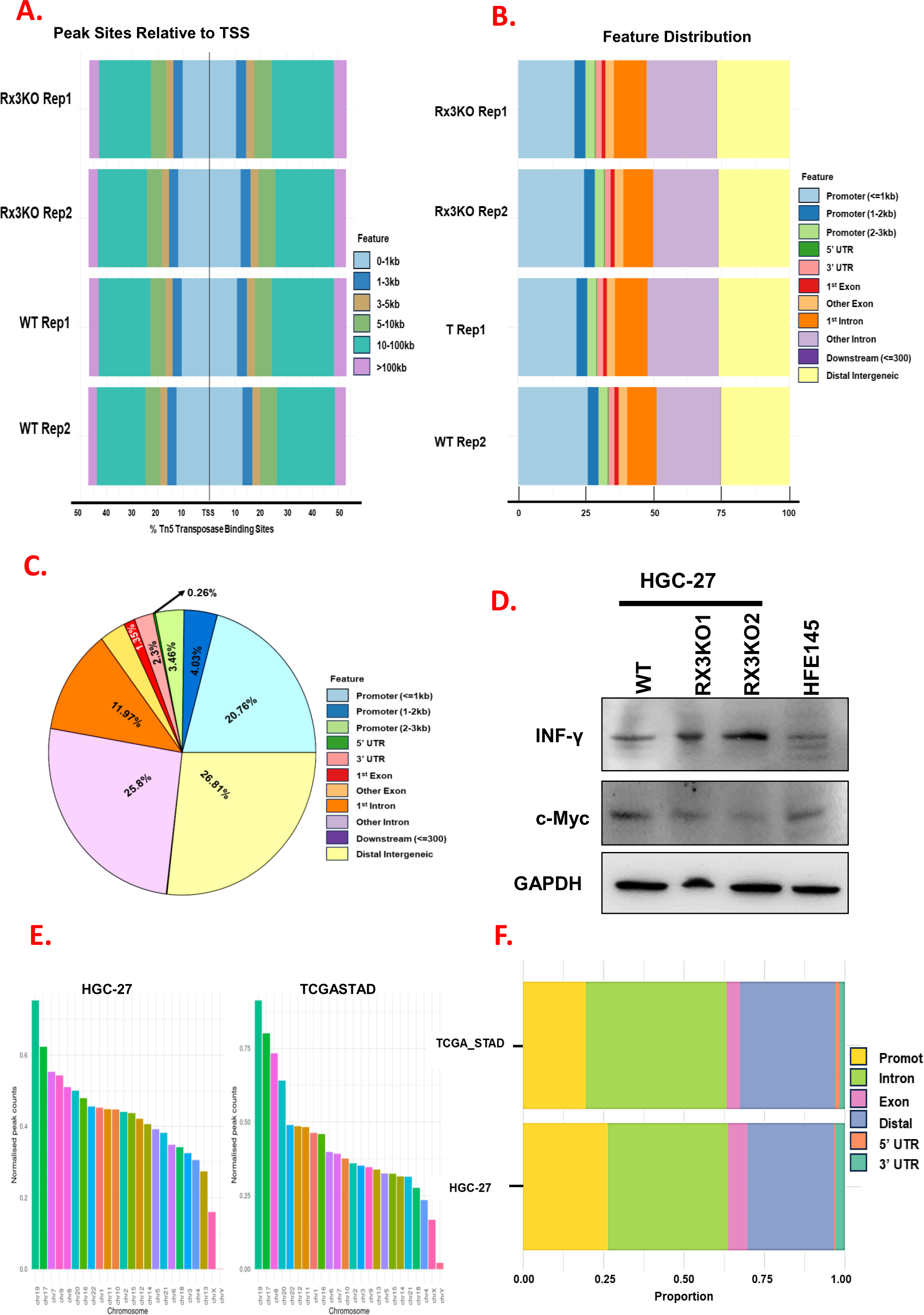
Chromatin Accessibility using ATAC-seq of Runx3 modulated expression cells and of the TCGA-STAD datasets. **A**). Representative profile plots of WT and Rx3KO show peaks enrichment and centered at the Transcription Start Site (TSS). **B).** Horizontal barplots of the genomic annotations across representative WT and Rx3KO samples showing distribution of overlapping genomic annotations (Left panel) and distribution of these peak sites around the TSS (right panel). **C).** Pie chart of genomic annotations percentages after using the default priority in the ChIPseeker annotatePeak function. **D).** Immunoblotting of represantive ATAC-seq and RNA seq dysregulated pathway targets. **E).** Normalized peak counts of Differentially Accessible (DA) regions in HGC-27 and The Cancer Genome Atlas Stomach Adenocarcinoma (TCGASTAD) dataset. **F).** Alignment of genomic annotations overlap between our HGC-27 and the TCGASTAD patient cohort ATAC-seq datasets.

**S3.**
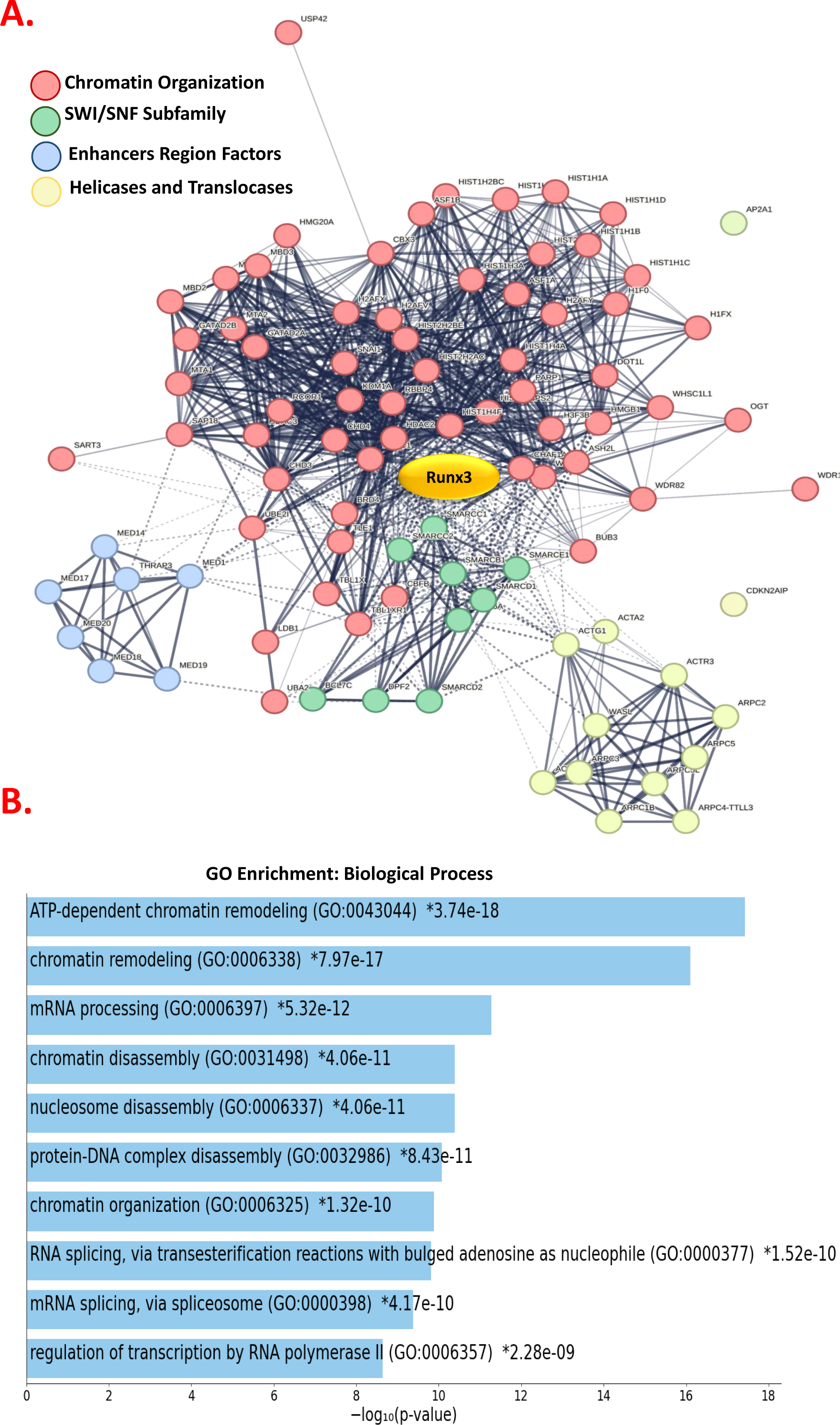
Runx3 interaction network in HGC-27 cells. **A)** Proteomics analysis of Runx3 interacting partners using STRING after Biotin-dependent proximity labelling followed by mass spectrometry analysis. **B).** The Gene Ontology (GO) annotations of enriched for biological processes.

**S4.**
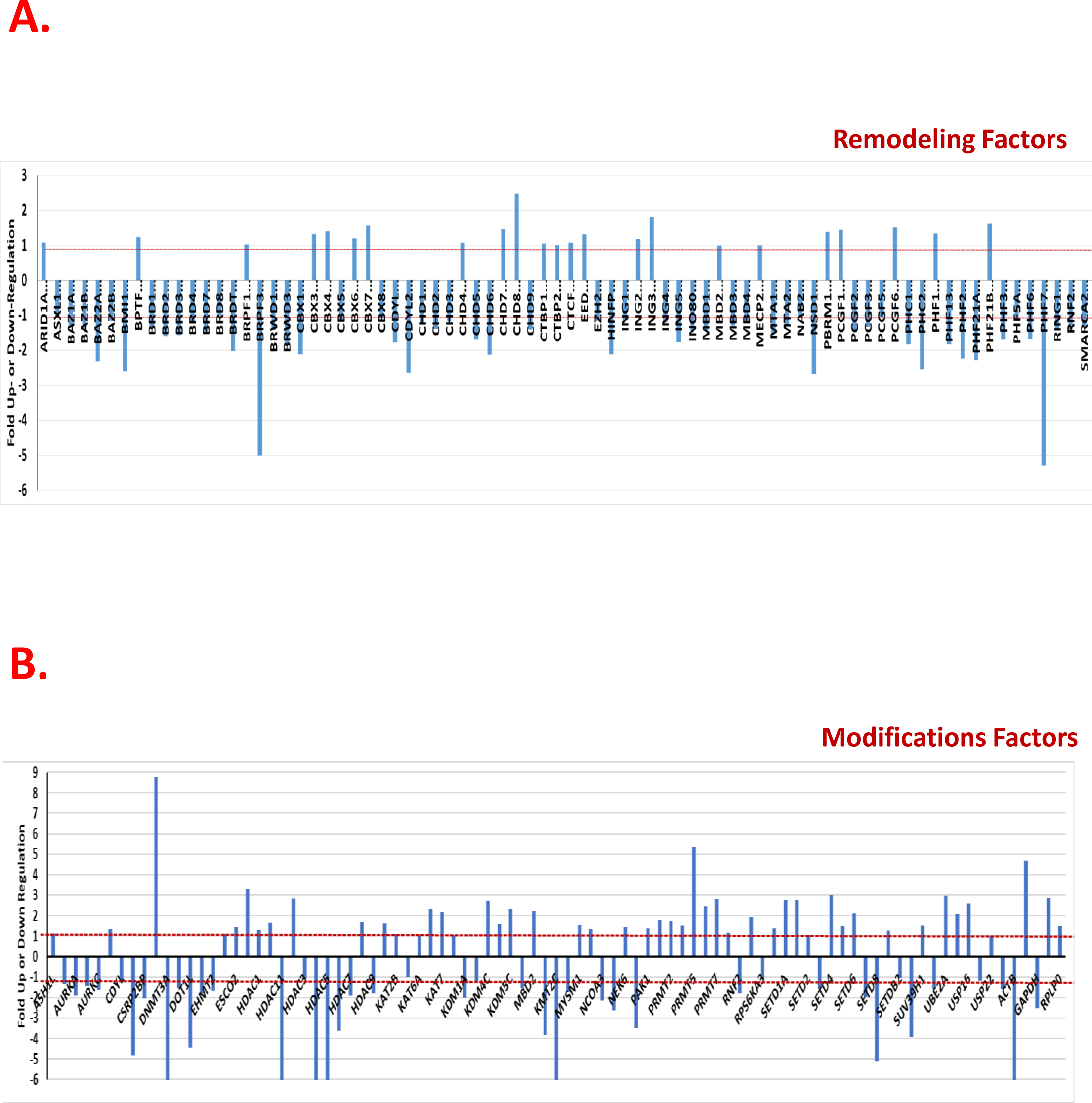
Runx3 ablation induced-gene expression changes using RT 2 Profiler PCR Arrays of human chromatin remoldeing and modification factors. Total RNA was extracted from WT and Rx3KO HGC-27 cells and analyzed by: **A).** Human chromatin remoldeing factors arrays (PAHS-086Z) and **B)**. Human chromatin modification enzymes arrays (PAHS-085Z). Fold changes >2.0 and a P-value < 0.05 were considered to be significant variations. The data represent the mean of three independent experiments.

**S5.**
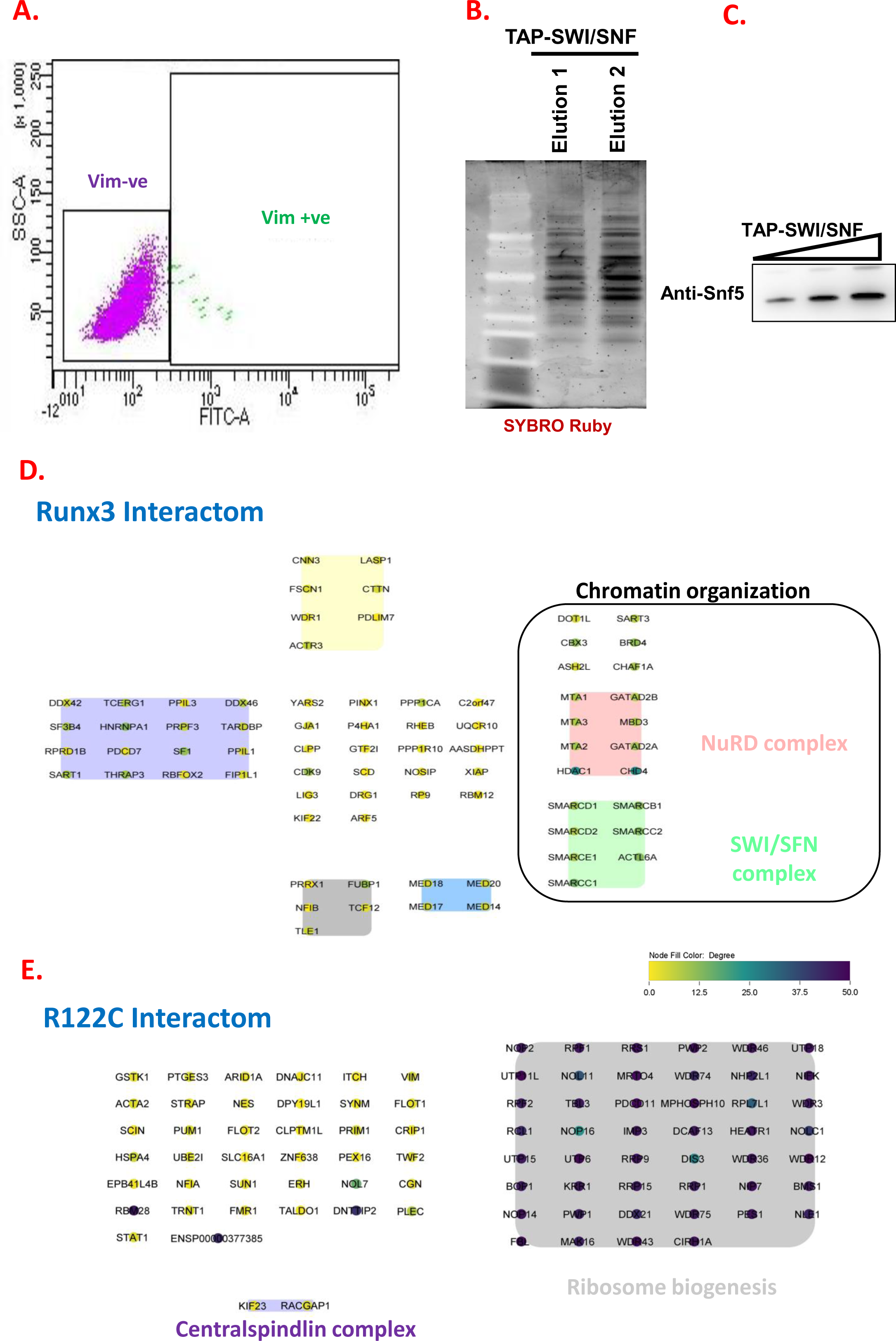
Runx3 interacting partners. **A).** Fluorescence Activated Cell Sorting (FCAS) sorting of SW13 vime –ve and vim+ve subtypes to establish correlation with SWI/SNF expression. **B).** Tandem Affinity Purification (TAP) of full length SWI/SNF complex from Euroscarf yeast TAP tagged strain followed by analysis of purity on 4-12% NuPAGE gels and SYBRO staining. **C).** Immunoblotting analysis of TAP-purified SWI/SNF using specific Snf5 antobody showing clear bands even at different titrated amounts (1-4μl). Interactom mapping of HGC-27 stably expressing: **D).** Dox-inducible 3×HA-TurboID-RUNX3 and the **E).** RUNT domain mutant R122C. Our data indicated that RUNT domain is cruicial for the interaction with chromatin factors specifically the SNF-2 subfamily as R122C loses interaction with SWI/SNF and NuRD components.

**S6.**
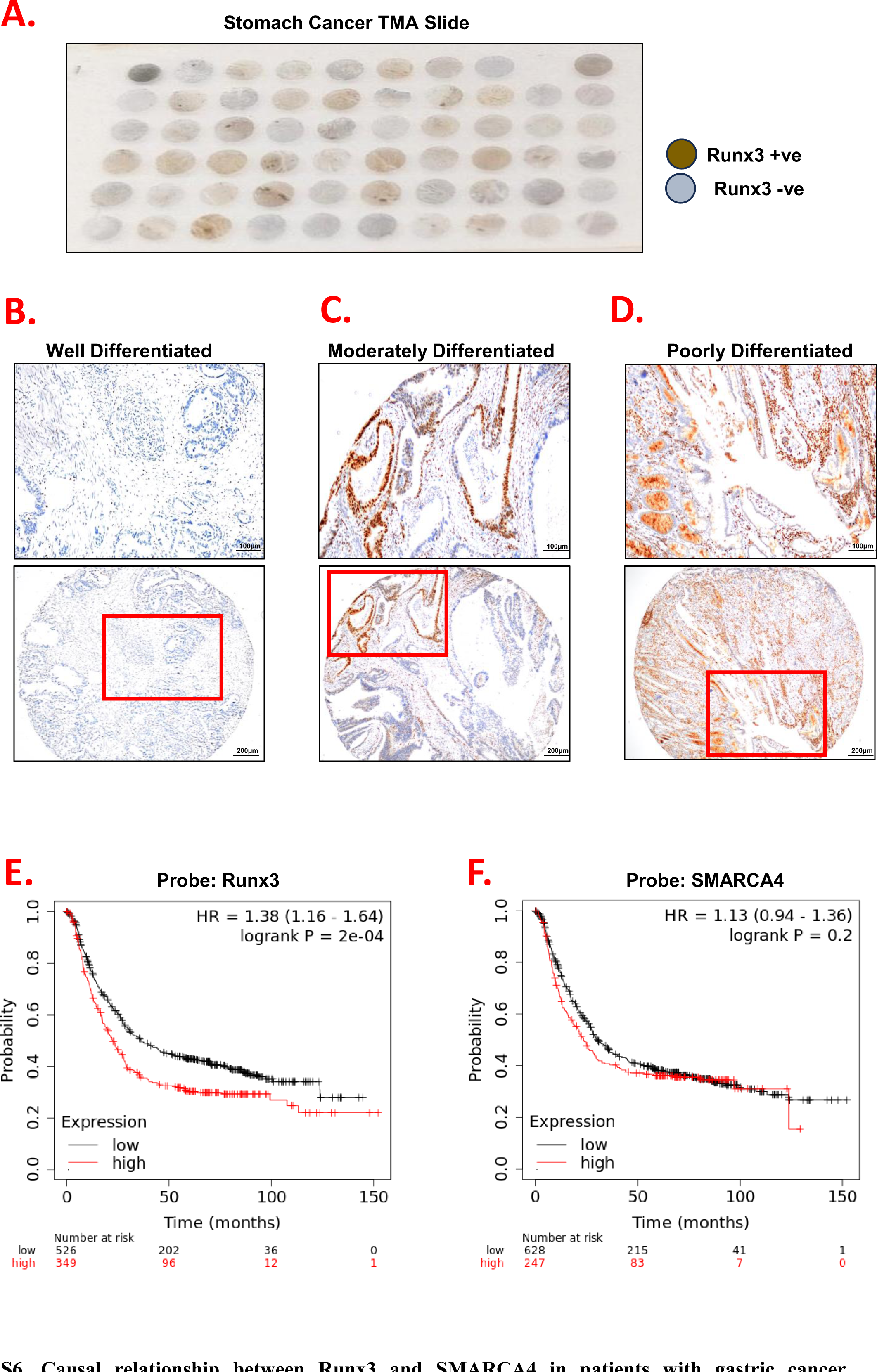
Causal relationship between Runx3 and SMARCA4 in patients with gastric cancer. Representative immunohistochemistry of Runx3 Tissue Array slide (n=59) of different gastric cancer tumor grades: **A).** Immunohistochemistry staining of whole Tissue Microarray (TMA) slide of different stages of gastric cancer from Novus Biologicals showing differential Runx3 expression depending on cancer stage and tissue anatomy. Kaplan–Meier curve depicting survival curve of: **B)**. Differentiated (Grade I, low grade), **C).** Moderately Differentiated (Grade II, intermediate), **D).** Poorly Differentiated (Grade III, high grade). Kaplan-Meier survival curve of gastric cancer patients (n=875) based on the following markers: **E).** Runx3, **F).** SMARCA4.

**S7.**
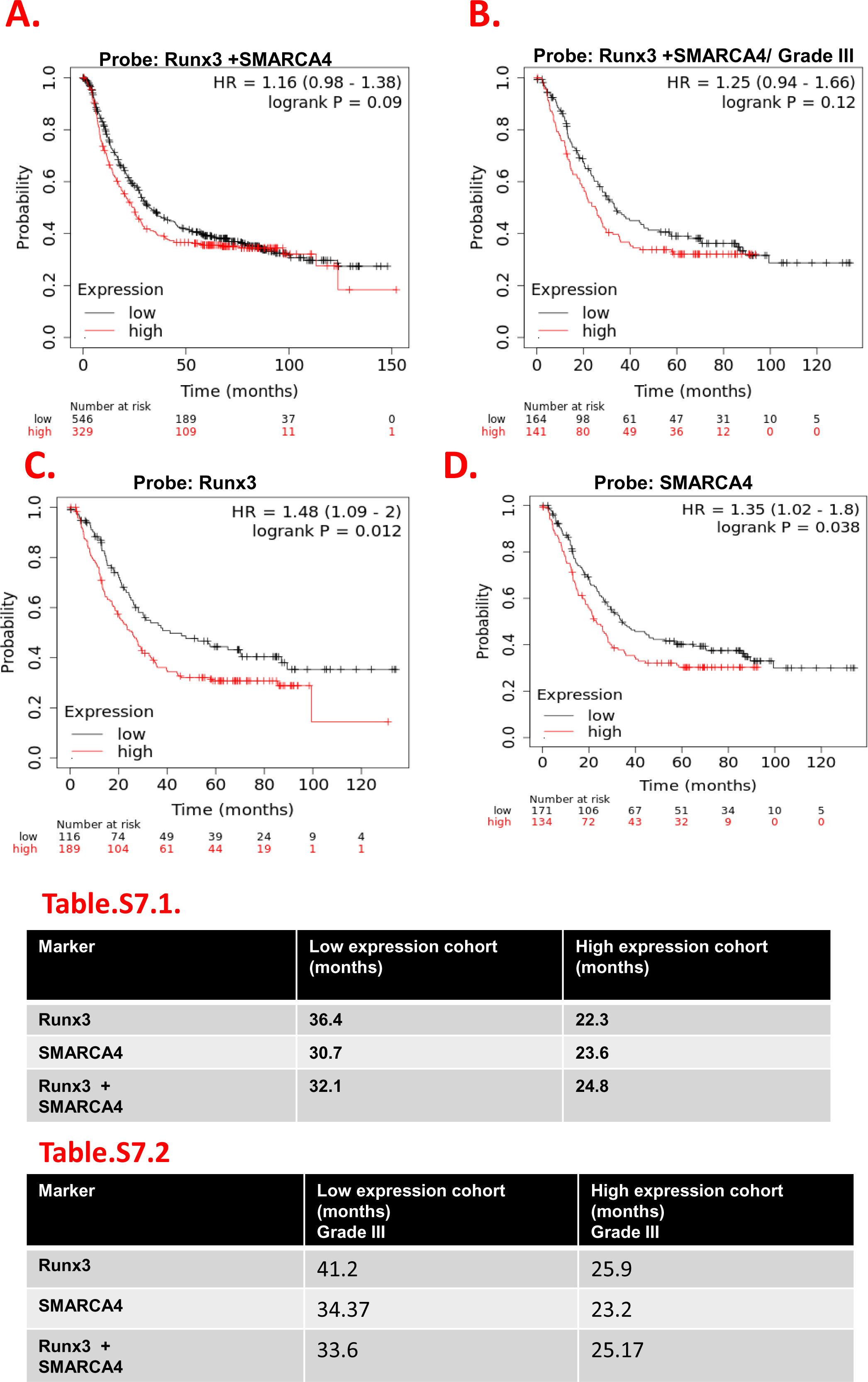
Runx3 expression correlation with SMARCA4 in gastric cancer patients. Kaplan-Meier survival curve of gastric cancer patients (n=875) based on the following markers: **A).** Combined markers of Runx3 and SMARCA4. **B).** Combined markers of Runx3 and SMARCA4 in cohort of Grade III cancer patients only (n=319). **C)** Overall gastric cancer cohort in patients with endogenous low Runx3 expression only and in **D).** Patients from Grade III only. **Tables summarizing median survival curve in: Table.S6.1.** Patients cohort with indicated markers’ (Runx3, SMARCA4 or combined) expression effect on prognosis. **Table.6S.2.** Based on patients with Grade III stomach cancer indicating better prognosis effect of Runx3, SMARCA4 or their combined downregulation.

**S8.**
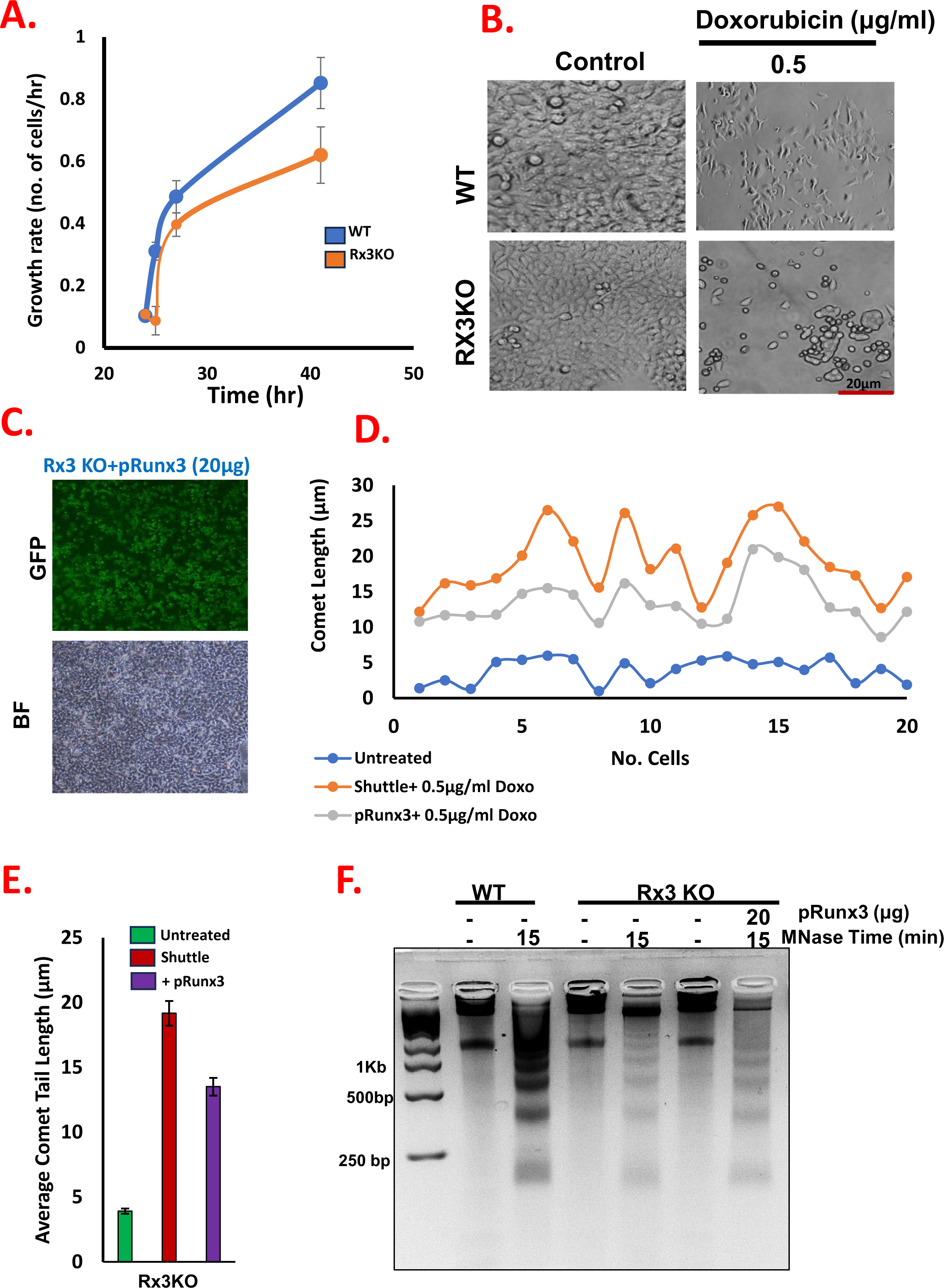
Runx3 depletion results in growth delay and DNA cellular defects in DNA damage repair restored after overexpression. **A).** Growth curves of HGC-27 cells showed decrease in Rx3KO viability and delayed growth after 30+hr compared to WT. **B).** Morphological changes observed in Rx3KO after DNA damage induction using 0.5μg/ml Doxorubicin (Doxo). Inverted phase contrast microscopy at 20X magnification images were obtained following Doxo treatment for 24hr. Elevated Runx3 expression in Rx3KO cells after 48hr post transfected with pRunx3 CRISPR resistant expression vector. **C).** Representative image of p-Runx3transfection efficiency in Rx3KO cells. **D).** Runx3 overexpression restores chromatin nucleosomal occupancy profile in Rx3KO cells. Increased mononucleosomes and decrease in higher polynucleoosmes were observedin Rx3KO cells after transfection with 20μg of pRunx3 followed by MNase digestion for 15min. **E).** Quantification of comet assay measured as relative comet length in 20 cells. Data are representative of three independent experiments. **F).** Average quantification of comet tail length.

