## Supplementary for "Runx3 Acts as Homodimeric Chromatin Binding Factor Regulating Heterochromatin-Mediated Cancerous Phenotype"

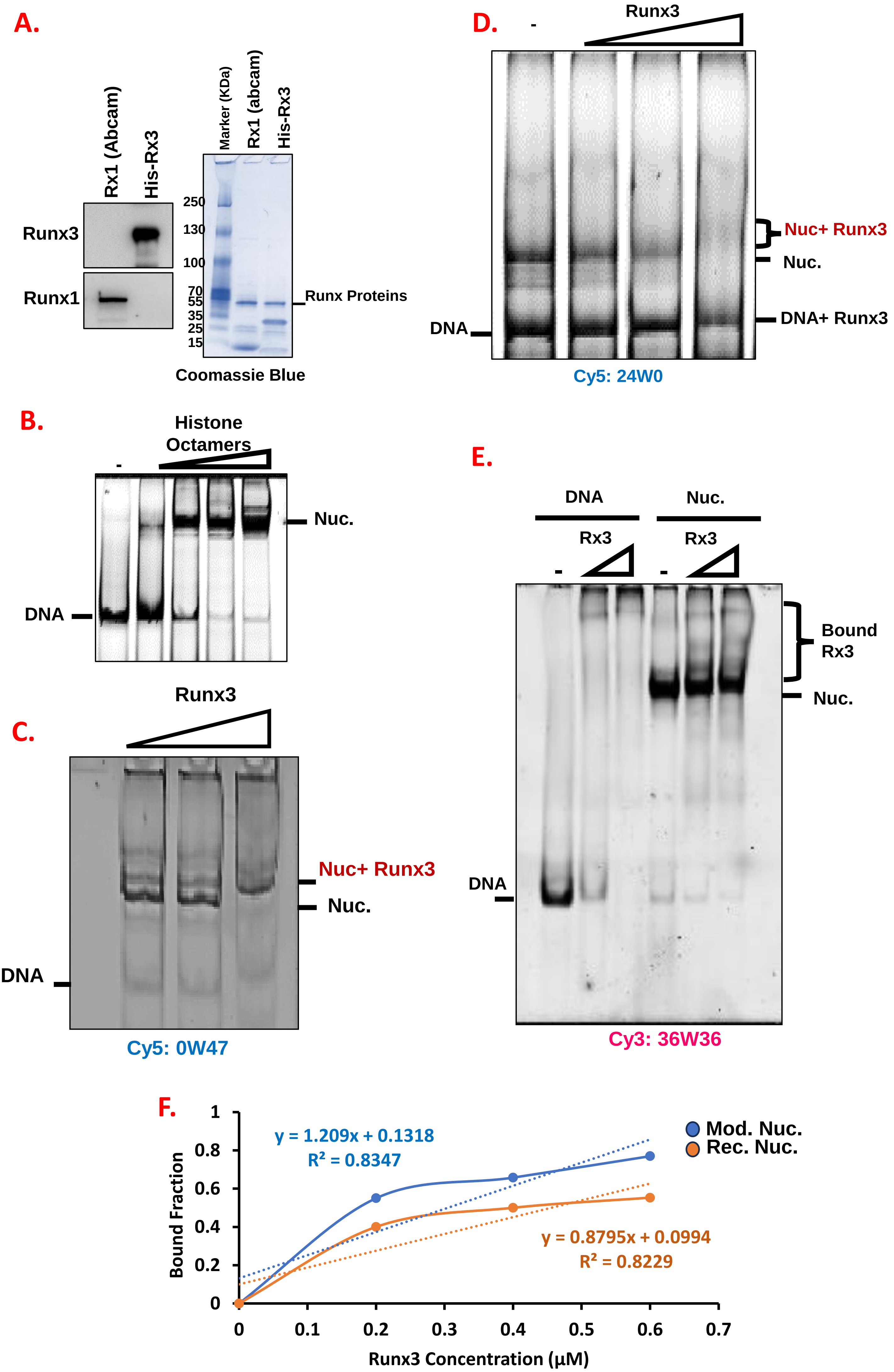


**S1. Runx3 binding to different nucleosomes structures and comparisons of affinity mangtitude. A).** Visualization of purified recombinant His-Runx3 and Runx1 (Abcam) by Commassie staining and Western Blotting using specific antibodies. **B).** Reconstitutions of fluorescent labelled mononucleoosmes at 1 µM concentration by stepwise dialysis. Comparisons of Runx3 binding affinity with: **C).** 0W47 mononucleosomes, **D).** 24W0 mononucleosomes. **E).** Binding of titrated equimolar amounts of Runx3 to DNA and 36W36 mononucleoosmes. In all reactions 30 nm of fluorescent labelled mononucleosomes were used. **F).** Quantification of Runx3 binding affinity to Cy-36W36 assembled into either modified nucleosomes (blue) or recombinant nucleosomes (orange).

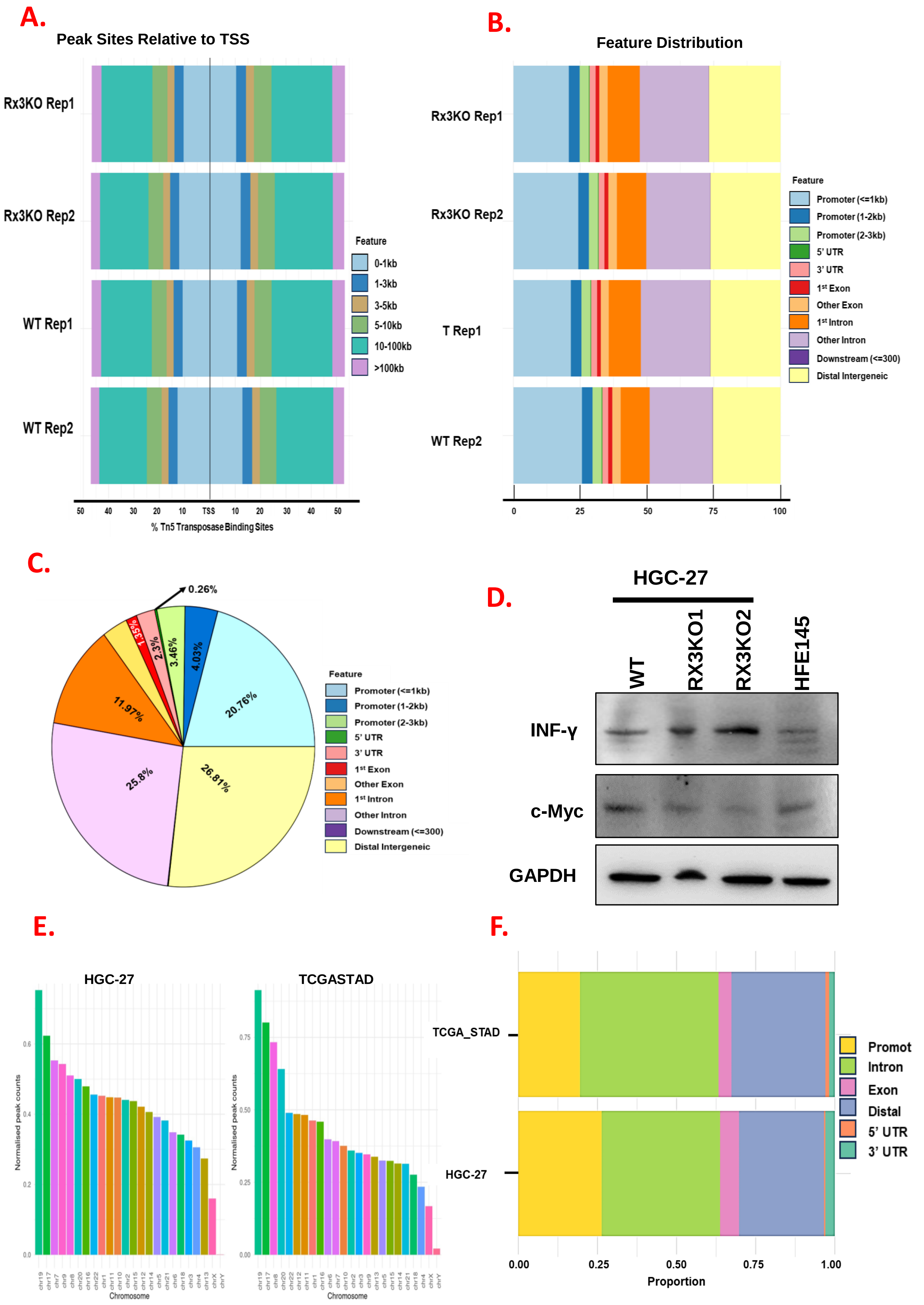


**S2. Chromatin Accessibility using ATAC-seq of Runx3 modulated expression cells and of the TCGA-STAD datasets. A**). Representative profile plots of WT and Rx3KO show peaks enrichment and centered at the Transcription Start Site (TSS). **B).** Horizontal barplots of the genomic annotations across representative WT and Rx3KO samples showing distribution of overlapping genomic annotations (Left panel ) and distribution of these peak sites around the TSS (right panel). **C).** Pie chart of genomic annotations percentages after using the default priority in the ChIPseeker annotatePeak function. **D).** Immunoblotting of represantive ATAC-seq and RNA seq dysregulated pathway targets. **E).** Normalized peak counts of Differentially Accessible (DA) regions in HGC-27 and The Cancer Genome Atlas Stomach Adenocarcinoma (TCGASTAD) dataset. **F).** Alignment of genomic annotations overlap between our HGC-27 and the TCGASTAD patient cohort ATAC-seq datasets.

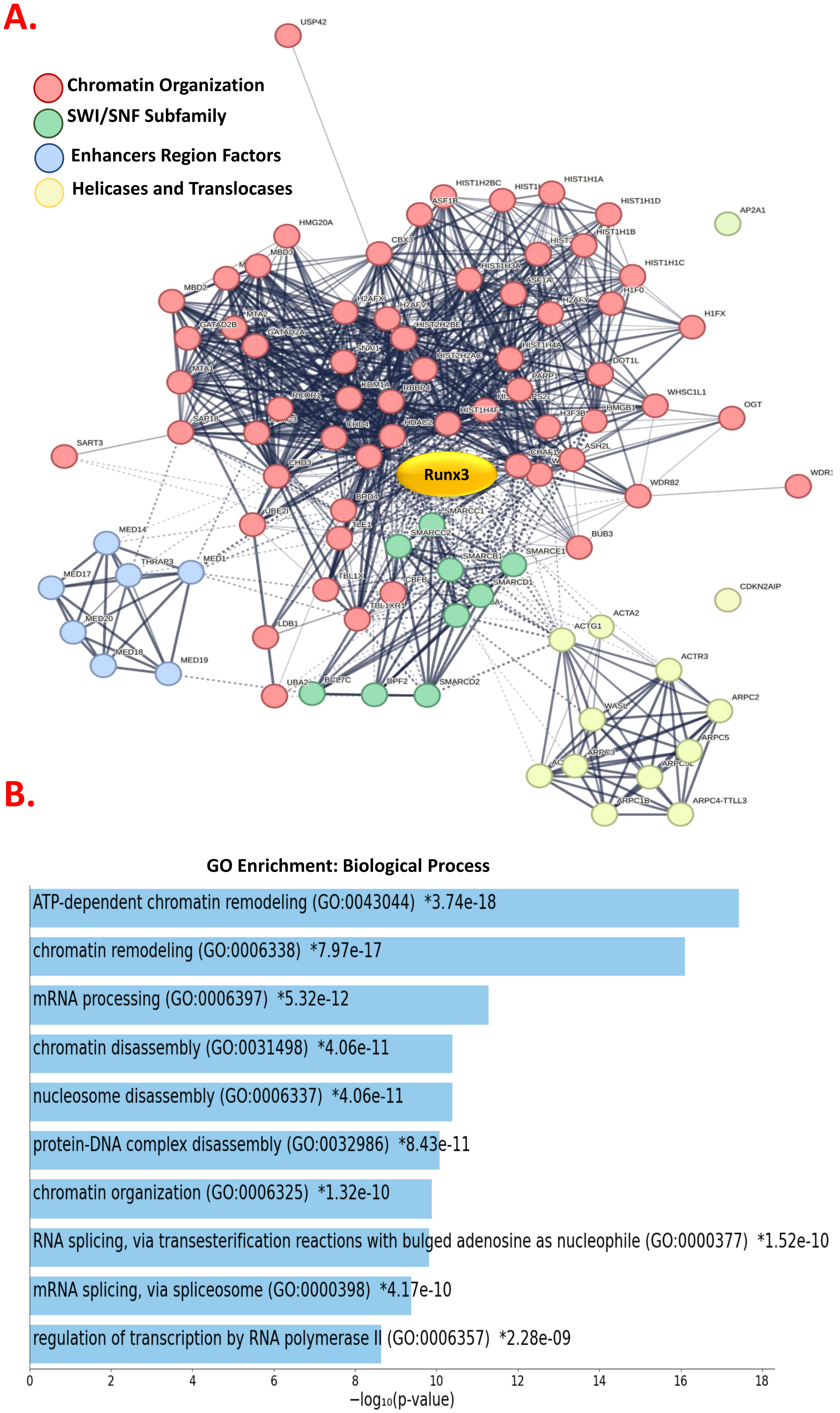


**S3. Runx3 interaction network in HGC-27 cells. A)** Proteomics analysis of Runx3 interacting partners using STRING after Biotin-dependent proximity labelling followed by mass spectrometry analysis. **B).** The Gene Ontology (GO) annotations of enriched for biological processes.

**A.**

Remodeling Factors

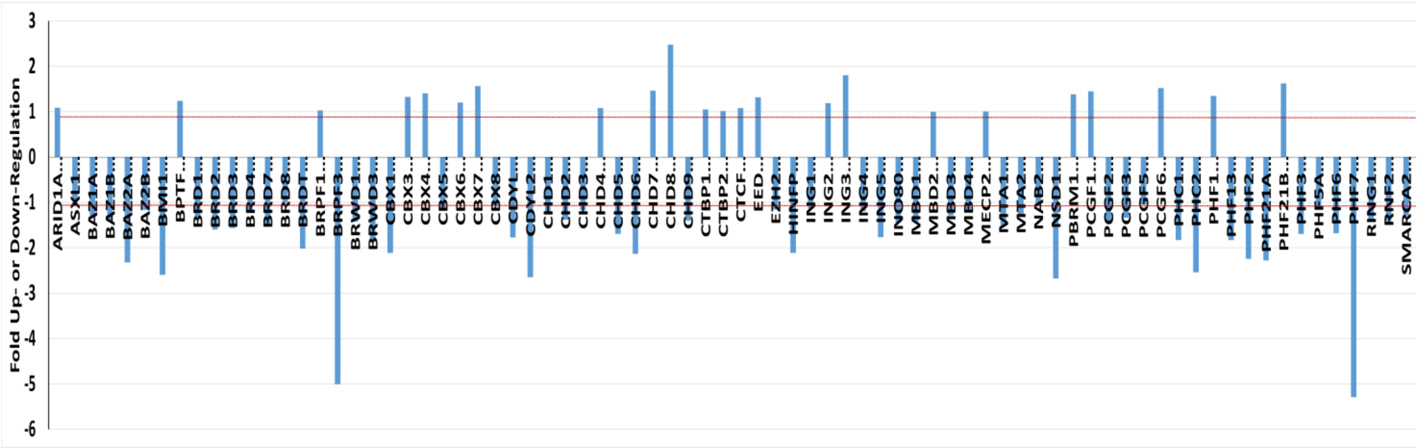


**B.**

Modifications Factors

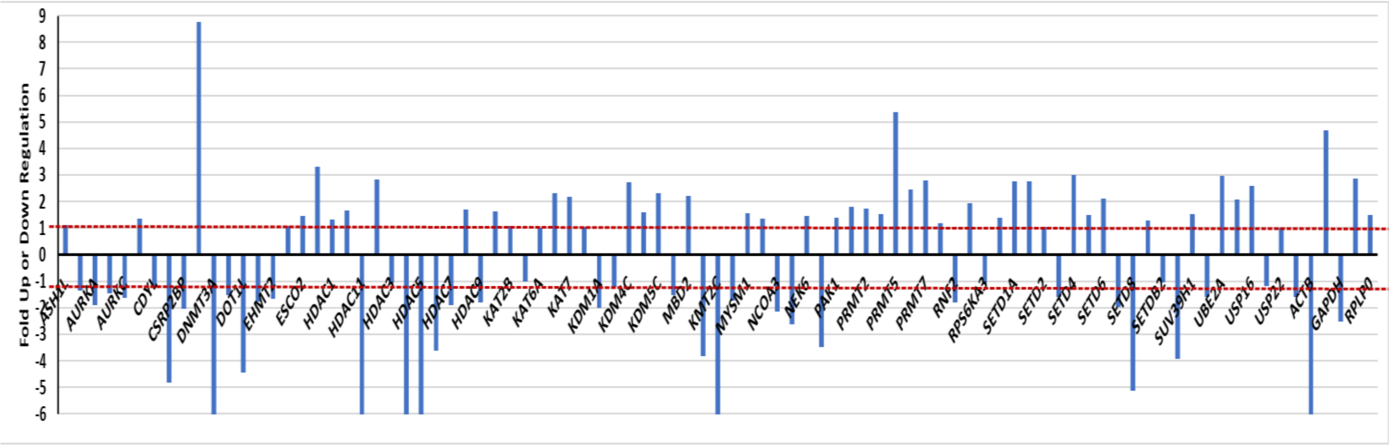


**S4. Runx3 ablation induced-gene expression changes using RT 2 Profiler PCR Arrays of human chromatin remoldeing and modification factors.** Total RNA was extracted from WT and Rx3KO HGC-27 cells and analyzed by: **A).** Human chromatin remoldeing factors arrays (PAHS-086Z) and **B)**. Human chromatin modification enzymes arrays (PAHS-085Z). Fold changes >2.0 and a P-value < 0.05 were considered to be significant variations. The data represent the mean of three independent experiments.

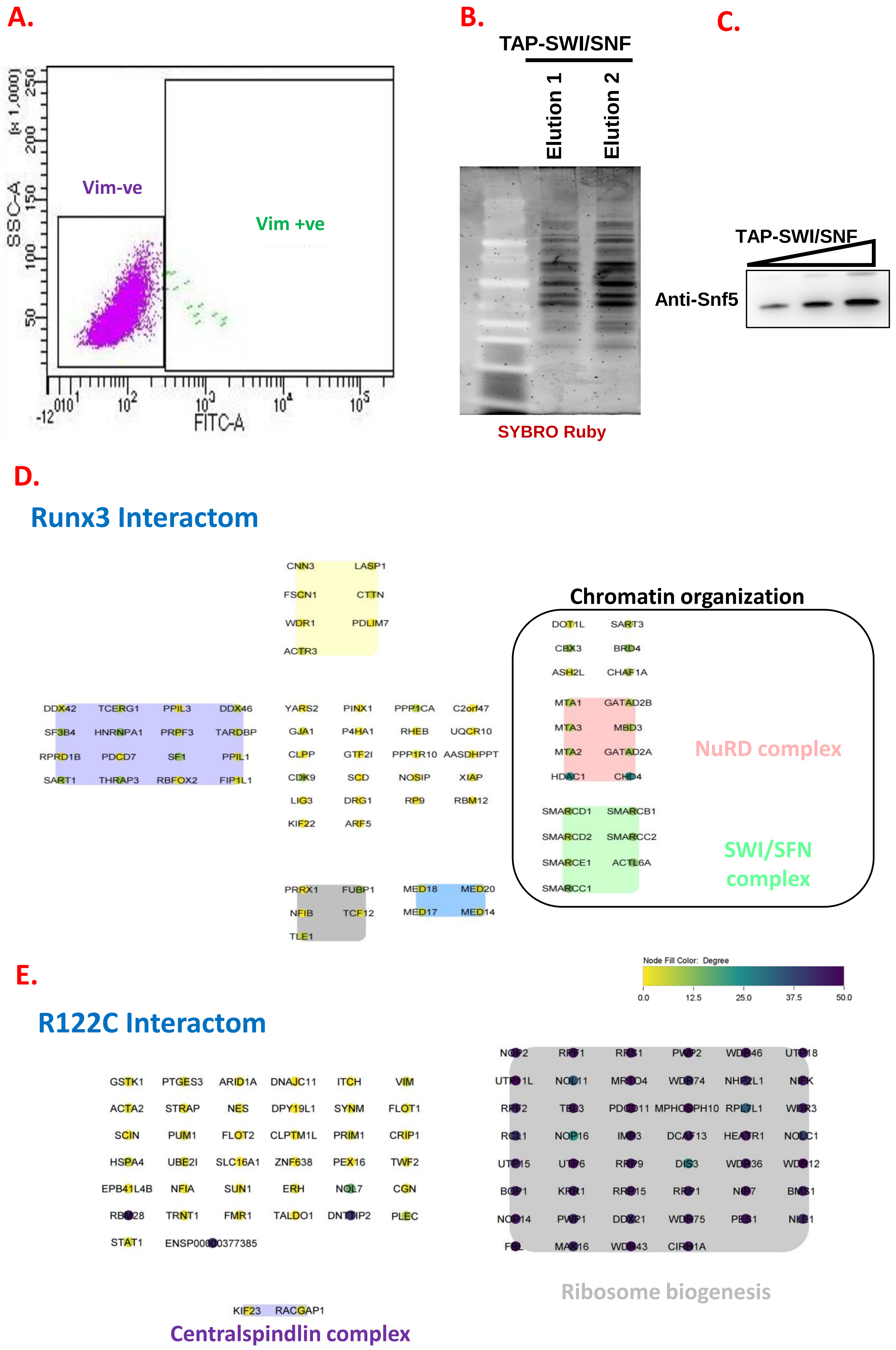


**S5. Runx3 interacting partners. A).** Fluorescence Activated Cell Sorting (FCAS) sorting of SW13 vime –ve and vim+ve subtypes to establish correlation with SWI/SNF expression. **B).** Tandem Affinity Purification (TAP) of full length SWI/SNF complex from Euroscarf yeast TAP tagged strain followed by analysis of purity on 4-12% NuPAGE gels and SYBRO staining. **C).** Immunoblotting analysis of TAP-purified SWI/SNF using specific Snf5 antobody showing clear bands even at different titrated amounts (1-4µl). Interactom mapping of HGC-27 stably expressing: **D).** Dox-inducible 3×HA-TurboID-RUNX3 and the **E).** RUNT domain mutant R122C. Our data indicated that RUNT domain is cruicial for the interaction with chromatin factors specifically the SNF-2 subfamily as R122C loses interaction with SWI/SNF and NuRD components.

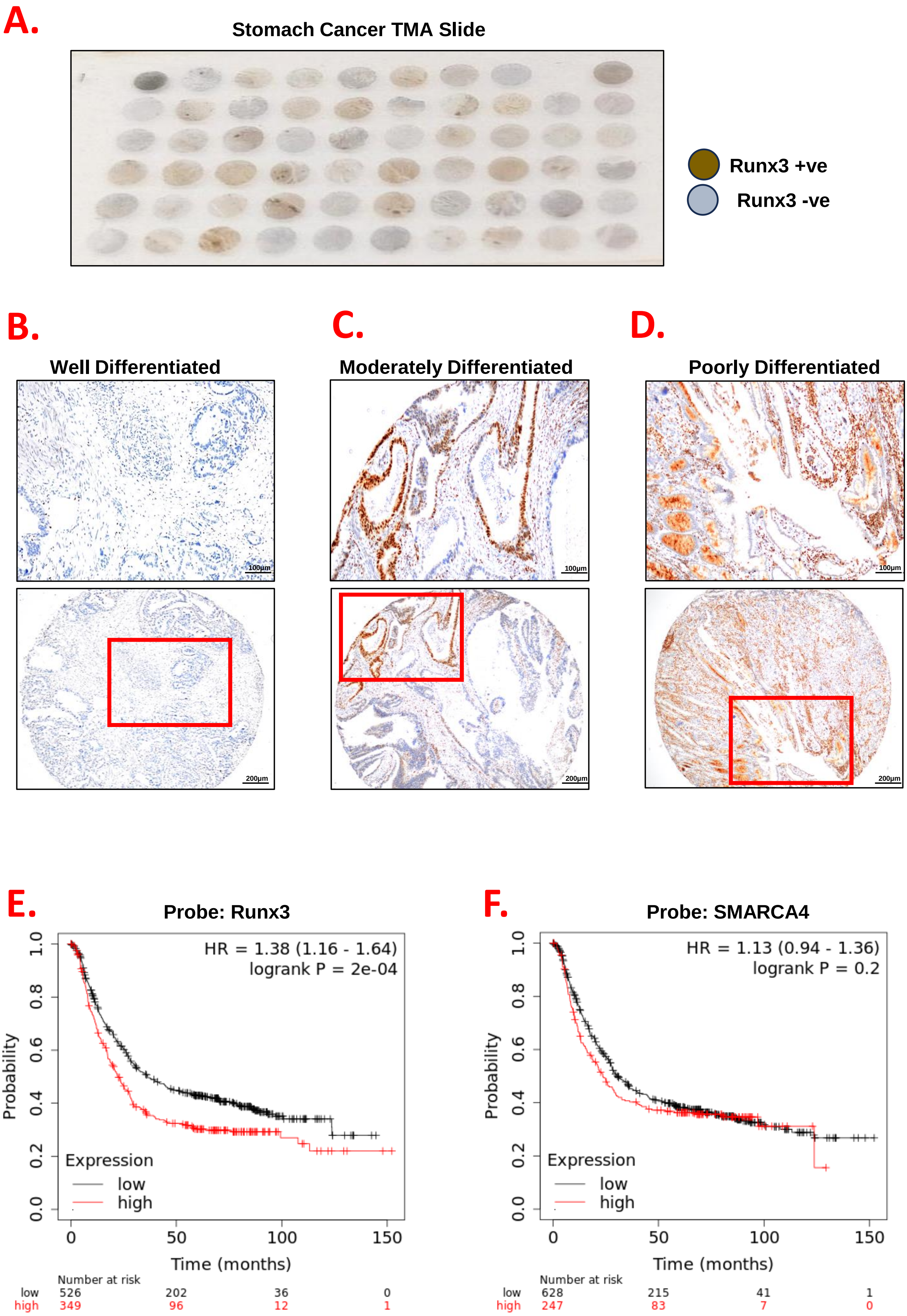


**S6. Causal relationship between Runx3 and SMARCA4 in patients with gastric cancer.** Representative immunohistochemistry of Runx3 Tissue Array slide (n=59) of different gastric cancer tumor grades: **A).** Immunohistochemistry staining of whole Tissue Microarray (TMA) slide of different stages of gastric cancer from Novus Biologicals showing differential Runx3 expression depending on cancer stage and tissue anatomy. Kaplan–Meier curve depicting survival curve of: **B).** Differentiated (Grade I, low grade), **C).** Moderately Differentiated (Grade II, intermediate), **D).** Poorly Differentiated (Grade III, high grade). Kaplan-Meier survival curve of gastric cancer patients (n=875) based on the following markers: **E).** Runx3, **F).** SMARCA4.

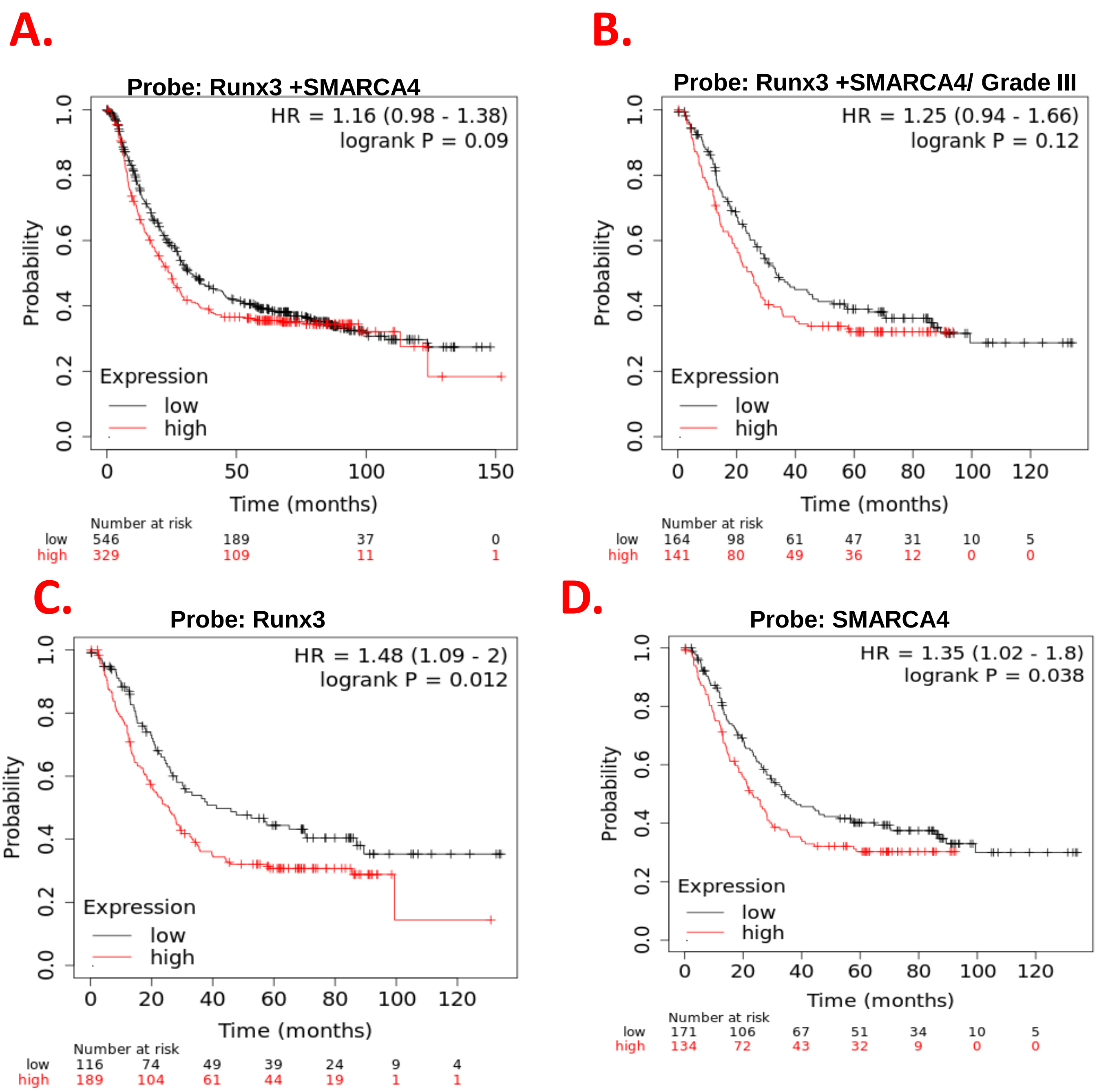


### Table.S7.1.

| Marker | Low expression cohort (months) | High expression cohort (months) |
|---|---|---|
| Runx3 | 36.4 | 22.3 |
| SMARCA4 | 30.7 | 23.6 |
| Runx3 + SMARCA4 | 32.1 | 24.8 |

### Table.S7.2

| Marker | Low expression cohort (months) Grade III | High expression cohort (months) Grade III |
|---|---|---|
| Runx3 | 41.2 | 25.9 |
| SMARCA4 | 34.37 | 23.2 |
| Runx3 + SMARCA4 | 33.6 | 25.17 |

**S7. Runx3 expression correlation with SMARCA4 in gastric cancer patients.** Kaplan-Meier survival curve of gastric cancer patients (n=875) based on the following markers: **A).** Combined markers of Runx3 and SMARCA4. **B).** Combined markers of Runx3 and SMARCA4 in cohort of Grade III cancer patients only (n=319). **C)** Overall gastric cancer cohort in patients with endogenous low Runx3 expression only and in **D).** Patients from Grade III only. **Tables summarizing median survival curve in: Table.S6.1.** Patients cohort with indicated markers' (Runx3, SMARCA4 or combined) expression effect on prognosis . **Table.6S.2.** Based on patients with Grade III stomach cancer indicating better prognosis effect of Runx3, SMARCA4 or their combined downregulation.

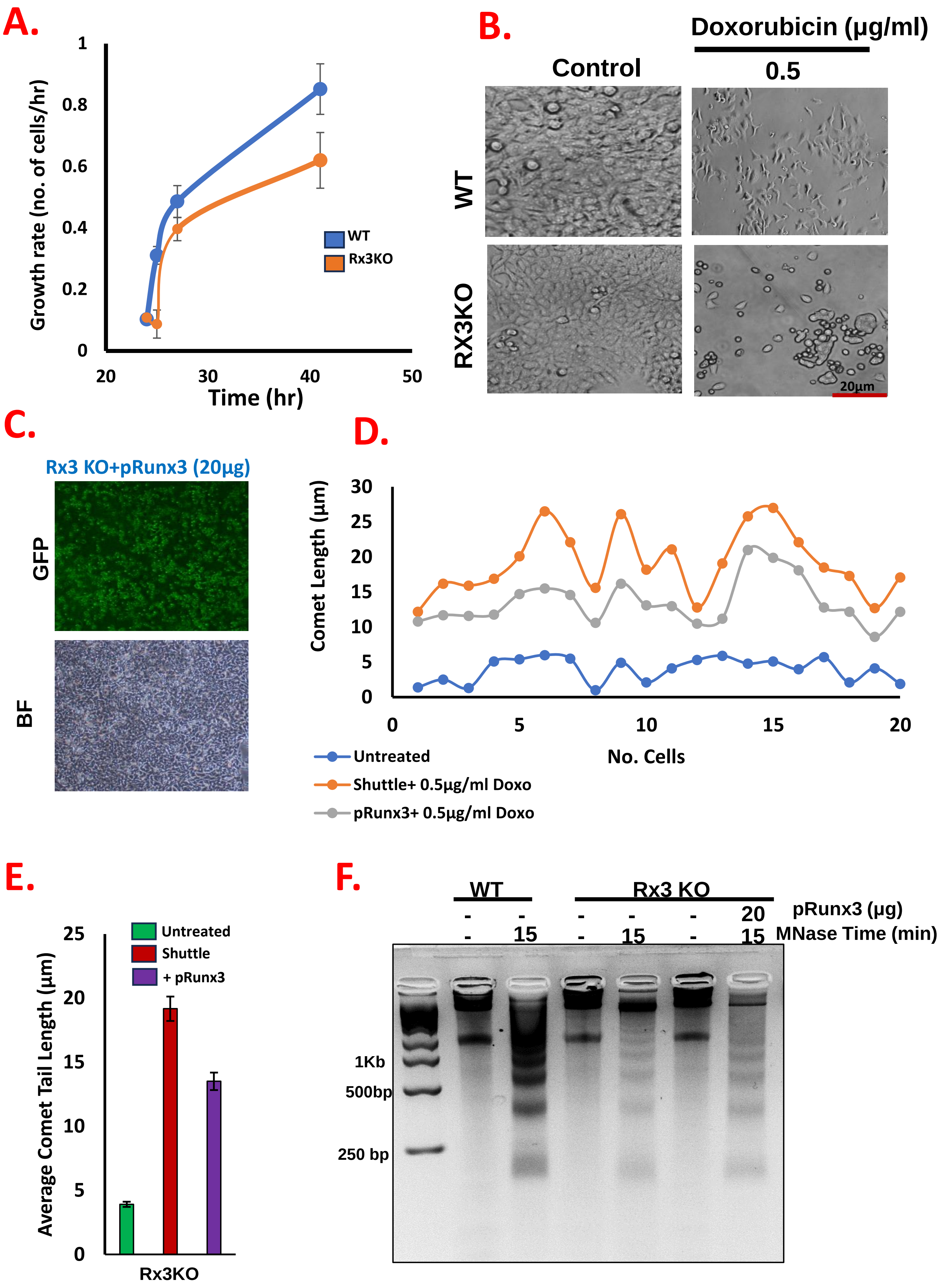


**S8. Runx3 depletion results in growth delay and DNA cellular defects in DNA damage repair restored after overexpression. A).** Growth curves of HGC-27 cells showed decrease in Rx3KO viability and delayed growth after 30+hr compared to WT. **B).** Morphological changes observed in Rx3KO after DNA damage induction using 0.5µg/ml Doxorubicin (Doxo). Inverted phase contrast microscopy at 20X magnification images were obtained following Doxo treatment for 24hr. Elevated Runx3 expression in Rx3KO cells after 48hr post transfected with pRunx3 CRISPR resistant expression vector. **C).** Representative image of p-Runx3transfection efficiency in Rx3KO cells. **D).** Runx3 overexpression restores chromatin nucleosomal occupancy profile in Rx3KO cells. Increased mononucleosomes and decrease in higher polynucleoosmes were observedin Rx3KO cells after transfection with 20µg of pRunx3 followed by MNase digestion for 15min. **E).** Quantification of comet assay measured as relative comet length in 20 cells. Data are representative of three independent experiments. **F).** Average quantification of comet tail length.
